# POWERDRESS-mediated histone deacetylation is essential for thermomorphogenesis in *Arabidopsis thaliana*

**DOI:** 10.1101/186916

**Authors:** Celine Tasset, Avilash Singh Yadav, Sridevi Sureshkumar, Rupali Singh, Lennard van der Woude, Maxim Nekrasov, David Tremethick, Martijn van Zanten, Sureshkumar Balasubramanian

## Abstract

Ambient temperature influences plant growth and development and minor changes can substantially impact crop yields. The underlying mechanisms for temperature perception and response are just beginning to emerge. Chromatin remodeling via the eviction of the histone variant H2A.Z in nucleosomes that alters gene expression is a critical component of thermal response in plants. However, whether chromatin-remodeling processes such as histone modifications play a global role in thermal response remains unknown. Using a combination of genetic analysis, chemical inhibition studies and RNA-seq analysis coupled with meta-analysis, here we identify POWERDRESS (PWR), a SANT-domain containing protein that is known to interact with HISTONE DEACETYLASE 9 (HDA9), as a novel key factor required for thermomorphogenesis in *Arabidopsis thaliana*. We identify that mutations in *PWR* impede thermomorphogenesis exemplified by severely attenuated temperature-induced hypocotyl/petiole elongation and early flowering. We show that inhibitors of histone deacetylases diminish temperature-induced hypocotyl elongation, which demonstrates for the first time a requirement for histone deacetylation in thermomorphogenesis. Genes that are misregulated in *pwr* mutants showed enrichment for GO terms associated with “response”. Our expression studies coupled with meta-analysis revealed a significant overlap between genes misregulated in *pwr* mutants and genes that are enriched for H2A.Z in their gene bodies. Meta-analyses reveal that genes misregulated in *pwr* mutants in diverse conditions also overlap with genes that are differentially expressed in the mutants of the components of the SWR1 complex that mediates H2A.Z nucleosome dynamics. Our findings thus uncover a role for PWR in facilitating thermal response and suggest a potential link between histone deacetylation and H2A.Z nucleosome dynamics in regulation of gene expression in plants.

**Author summary:** Plant growth and development is influenced by a variety of external environmental cues. Ambient temperature affects almost all stages of plant development but the underlying molecular mechanisms remain largely unknown. In this paper, the authors show that histone deacetylation, one of the major chromatin remodeling processes, is essential for eliciting growth temperature-induced responses in plants. The authors identify POWERDRESS, a protein known to interact with HISTONE DEACETYLASE 9, as a novel key player essential for eliciting high temperature induced responses in Arabidopsis. Another chromatin remodeling mechanism that is known to play a role in thermal response is the eviction of histone variant H2A.Z from nucleosomes. Through transcriptome studies the authors demonstrate an overlap between gene regulations conferred through PWR-mediated histone H3 deacetylation and that conferred via histone H2A.Z eviction/incorporation dynamics. This study identifies a key novel gene that is essential for plants to elicit high temperature responses and reveals close links between two seemingly distinct chromatin-remodeling processes in regulating gene expression in plants.

## Introduction

Plant development is highly sensitive to their growth environment. Ambient temperature, one of the major environmental factors that influences plant growth has a significant impact throughout plant development [1]. Even minor changes in temperature can modulate life history traits such as flowering time and seed set [2, 3]. Elevated temperatures modulate plant growth and development in a process termed “thermomorphogenesis” that results in a suite of phenotypes including an increase in hypocotyl elongation, petiole elongation and early flowering [1]. This is also often coupled with a dampening of defense response [4]. The molecular basis of thermal response is just beginning to emerge and appear to involve primarily changes at the level of transcription [1]. Recent work suggests that thermal cues are in part perceived through the photoreceptor phytochromes [5, 6]. For example phyB has been shown to bind to the promoters of its target genes in a temperature-dependent manner modulating transcriptional response to temperature [5].

One of the other key molecular events in thermal response involves chromatin remodeling associated with the eviction/incorporation dynamics of the histone variant H2A.Z in the nucleosomes [7]. Higher temperatures has been shown to lead to the eviction of the histone variant H2A.Z from nucleosomes [7]. H2A.Z eviction upon exposure to elevated temperature loosens the chromatin resulting in changes in gene expression [7]. H2A.Z nucleosome dynamics appears to be critical not only for temperature response, but also for general response to external stimuli [8]. H2A.Z is enriched in gene bodies of the “responsive genes”, and *h2a.z* mutants display mis-regulation of genes associated with response to environmental stimuli [8]. The eviction/incorporation of H2A.Z on to the nucleosomes is mediated through the SWR1 complex in Arabidopsis that consists of proteins encoded by *ACTIN RELATED PROTEIN 6 (ARP6), SWC6* and *PHOTOPERIOD INDEPENDENT EARLY FLOWERING 1 (PIE1).* Mutations in these genes result in pleiotropic phenotypes [9-12]. In contrast to our understanding of the temperature-induced H2A.Z eviction, very little is known about the global role of other chromatin remodeling processes such as histone modifications in ambient temperature response [1, 13].

A central integrator in this transcriptional network is *PHYTOCHROME INTERACTING FACTOR 4 (PIF4)*, which encodes a transcription factor that mediates several temperature-associated phenotypes including defense response [1, 14-18]. PIF4 is regulated at multiple levels via complex transcriptional and post-transcriptional regulatory mechanisms in response to temperature [1]. Subsequently, PIF4 regulates downstream target genes, primarily through transcription [1]. For example, PIF4 regulates temperature-induced hypocotyl elongation via stimulating auxin biosynthesis by binding to the promoters of auxin biosynthesis genes, including *YUCCA8* [14-16, 19]. Thus transcriptional responses at multiple levels play critical roles in governing temperature responses in plants.

Here, we identify POWERDRESS (PWR), which is known to interact with HISTONE DEACETYLASE 9 (HDA9) [20, 21] as a novel key factor that is essential for thermomorphogenesis in *Arabidopsis*, and uncover a central role for histone deacetylation in mediating thermal response. We demonstrate that blocking histone deacetylation abolishes temperature-induced hypocotyl elongation. Through RNA-seq experiments, we show that *PWR* suppresses defense gene expression at elevated temperatures. Using our RNA-seq data with the meta analysis of published genome-wide H2A.Z ChIP-seq data we show that there is a significant overlap between genes that are mis-regulated in the *pwr* mutants and those that are modulated through H2A.Z eviction/incorporation dynamics. Thus, our findings reveal a global role for histone deacetylation in thermal response. In addition our findings also show a general association between two distinct chromatin remodeling mechanisms *viz.,* histone H3 deacetylation and H2A.Z nucleosome dynamics in regulating gene expression that extends beyond thermal response.

## Results

Elevated temperatures result in increased hypocotyl elongation in *Arabidopsis thaliana* [19]. To identify new genes that facilitate thermomorphogenesis, we carried out a forward genetic screen. T-DNA insertion lines [22] allow simultaneous screening of phenotypes at multiple conditions. We screened more than 7000 lines at 23 °C and 27 °C for attenuated response in temperature-induced hypocotyl elongation and identified 4 potential mutants with altered thermal response. One of the lines that carried an insertion at At3g52250/*POWERDRESS (PWR)* displayed a severely diminished thermal response in hypocotyl elongation (Fig. 1A, B, *pGxE<0.0001*). We examined additional independent T-DNA insertion lines at this locus and found the reduction-of-function lines to display a reduced thermal response, which suggests that *PWR* is essential for temperature-induced hypocotyl elongation (Fig. 1B & S1). An *ems-*induced mutant for *PWR (pwr-1),* has been previously isolated as an enhancer of *agamous* [23]. This *pwr-1* allele in the L*er* background also displayed a similar impairment in temperature-induced hypocotyl elongation (Fig 1B, *pGxE<0.0001*). The impaired thermal response was abolished in the *pPWR::PWR-GFP* line, independently confirming that *PWR* is the causal locus for the attenuated thermal response (Fig. 1B).

**Fig. 1.**
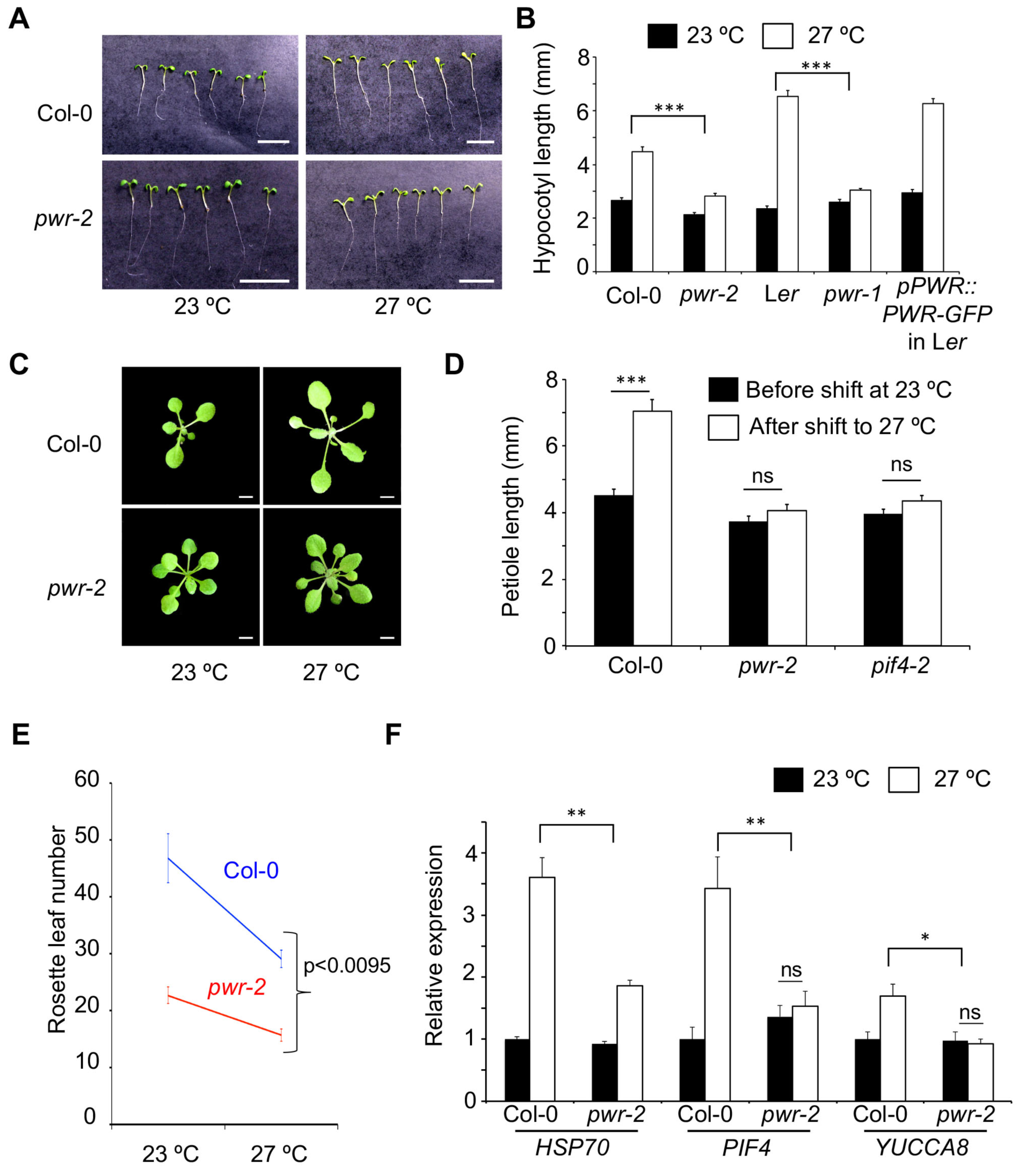
Mutations in *PWR* attenuate thermal responses. A) Hypocotyl lengths of *pwr-2* compared to Col-0 at two different temperatures in short days. B) Hypocotyl length of various genotypes at different temperatures. The *p-value* for the GxE interactions are shown above the bar graphs. *pwr-2* is in the Col-0 background. *pwr-1* and *pPWR::PWR-GFP* are in the L*er* background. N=15 for all samples. C) Petiole elongation of Col-0 and *pwr-2* two days after shift to 27 °C. The same plants are shown before and after the shift. D) Quantification of petiole elongation. N=23‐ 45. E) Flowering time measured as rosette leaf number in Col-0 and *pwr-2* at 23 °C and 27 °C with *p-valu*e for GxE interaction. F) Relative expression levels of *HSP70, PIF4* and *YUCCA8* in Col-0 and *pwr-2*, in 2-week old seedlings at 23 °C and 27 °C. Data are averages from three independent biological replicates, with each representing approximately 25-30 seedlings. *p-value* for the GxE interaction is shown. *p-values*: ***<0.0001, **<0.001, *<0.05, ns=not significant. Error bars represent standard error. Scale bars in A and C represent 5mm.

To assess whether the *pwr* mutation specifically affects hypocotyl development or generally impairs thermomorphogenesis, we evaluated other temperature-associated phenotypes. Elevated temperatures increase petiole length [14] (Fig. S2A) and plants, when shifted from 23 °C to 27 °C display an elongated petiole within 2 days (Fig. 1C-D). This marked response to temperature-shift was not observed in *pwr-2* mutants (Fig. 1C-D). Higher temperatures result in early flowering in Arabidopsis [3]. While mutations in *pwr* also result in early flowering [23] (Fig. S2B), the thermo-sensitivity of floral induction was significantly reduced in *pwr-2* (Fig. 1E, *pGxE<0.0095*). In addition *pwr-2* mutants appeared smaller than wild type plants (Fig. S2B), which suggests that there is a general impairment of plant growth in *pwr-2*. The observed reduction in temperature-sensitivity correlated with an attenuated response in the temperature-induced expression of *HSP70, PIF4* and *YUCCA8* (Fig. 1F), genes known to be induced upon elevated temperatures. Taken together these results suggest that *pwr* mutants are generally impaired in thermomorphogenesis.

The attenuated expression of *PIF4* and *YUCCA8* in *pwr-2* (Fig. 1F) suggests that *PWR* is required for the temperature-induced *PIF4* expression and subsequent auxin biosynthesis, which could be the underlying mechanism for the impaired thermal response in hypocotyl elongation. In contrast, *PWR* expression remained mostly unaltered in the *pif4-2* mutants (Fig. S3), which suggests that *PIF4* does not regulate *PWR* at the transcriptional level. Consistent with the idea that *PWR* and *PIF4* act in the same genetic cascade, the *pif4 pwr* double mutants were not significantly different from either single mutant, *i.e.,* no additive or antagonistic interactions were observed in temperature-induced hypocotyl elongation (Fig 2A, Fig. S4A). However, *pwr* and *pif4* mutants significantly differ in their flowering phenotype both at 23 °C and 27 °C with the *pwr* mutants displaying early flowering at both temperatures [23-25] (Fig. 1E,). We found *pwr-2 pif4-2 or pwr-2 pif4-101* double mutants to be early flowering similar to *pwr-2* mutants (Fig 2B, Fig. S4B). The early flowering at 27 °C was associated with an increase in *FT* expression, which suggests that the loss of *PWR* can overcome the requirement for the proposed activation of *FT* expression by *PIF4*[24] at high ambient temperatures (Fig. 2C).

**Fig. 2.**
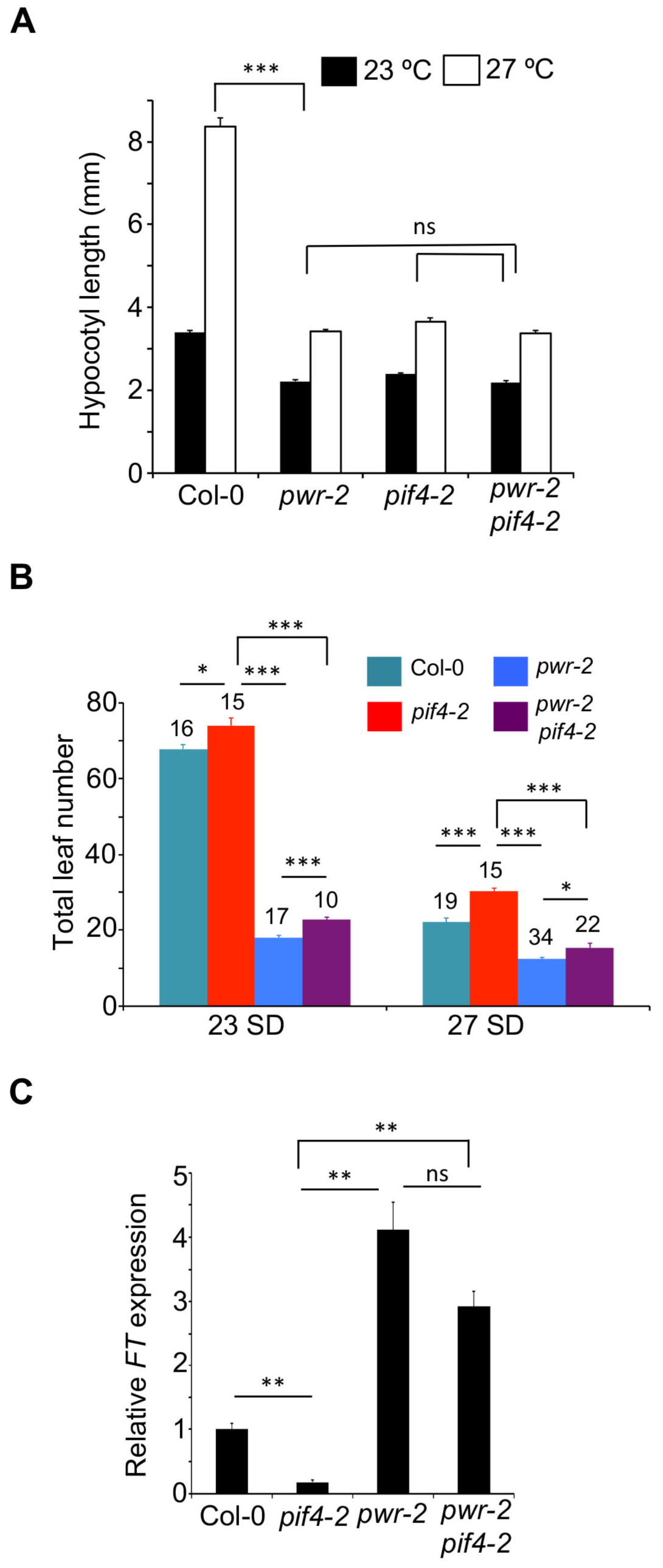
*PWR* acts upstream of *PIF4.* A) Hypocotyl lengths of various genotypes at 23 °C and 27 °C*. p-values* for the corresponding GxE interactions determined through ANOVA are shown. B) Flowering time measured as total leaf number in *pif4-2 pwr-2* double mutants compared to single mutants at two different temperatures in short days (please see Supplementary Figure 4 for the data with *pif4-101)*. Number of plants and the *p-values* determined by Student’s t-test are shown above bar graphs. C) Relative *FT* expression levels in different genotypes at 27 °C. The data is normalized against the Col-0 reference with *TUBULIN* as internal control. Error bars represent standard error. *p-values*:***<0.0001, **<0.001, *<0.05, ns=not significant.

PWR contains a SANT domain that has been suggested to play role in regulating chromatin accessibility by mediating the interaction between histone tails and the histone modifying enzymes [26]. To assess whether the acetylation status of histones modulate temperature-induced hypocotyl elongation, we grew plants in presence of histone acetylation/deacetylation inhibitors. While we did not detect any difference in hypocotyl length presence of histone acetylation inhibitor curcumin (Fig. S5), temperature-induced hypocotyl elongation was severely compromised in plants grown in presence of different histone deacetylase (HDAC) inhibitors *viz.,* sodium butyrate, Droxinostat, CP64434 hydrate or trichostatin A (Fig. 3A, S6A & S6B). Western blots confirmed inhibition of deacetylation with an increase in acetylated proteins in presence of sodium butyrate (Fig. S6C). These findings confirmed that histone deacetylation is essential for thermomorphogenesis. Comparison of *pwr-2* mutants with wild type Col-0 revealed significant drug x temperature (Fig. 3B, S6B) and drug x genotype (Fig. S6D) interactions confirming that the effect of histone deacetylation depends on the genotype and temperature. The effect of HDAC inhibitors were less pronounced in *pwr-2* compared to Col-0, suggesting that *PWR* acts in the same pathway that is targeted by the HDAC inhibitors. Correlating with the phenotypes, there was a reduction in temperature-induced expression of *YUCCA8* and *PIF4* in presence of HDAC inhibitors (Fig. 3C, S6E). Moreover, *hda9* mutants also displayed attenuated responses in temperature-induced hypocotyl elongation (Fig. 3D). Consistent with the recent findings [20, 21], we detected an increase in H3K9-acetylation in *pwr-2* mutants (Fig. 3E). In addition, we also detected an increase in acetylated H2A.Z in *pwr-2* (Fig. 3E). Taken together these data suggests that PWR-mediated histone deacetylation is essential for thermomorphogenesis in Arabidopsis.

**Fig. 3.**
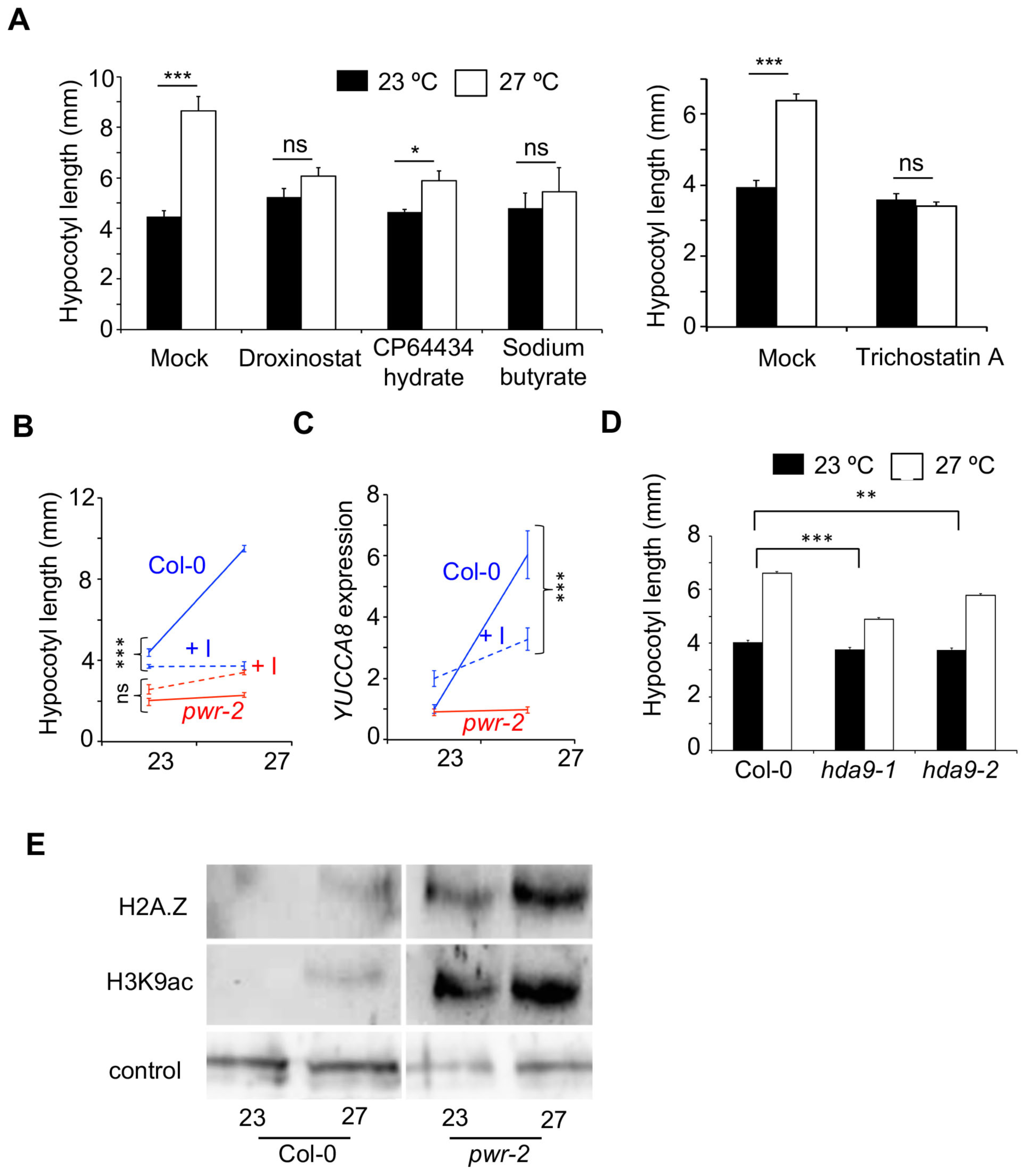
Histone deacetylation is essential for temperature-induced hypocotyl elongation. A) Hypocotyl lengths of Col-0 plants grown in presence of different deacetylase inhibitors (at 10uM except for Trichostatin A (1uM) and a mock control at 23 °C and 27 °C. *p-values* for differences in hypocotyl elongation between 23 °C and 27 °C determined by a Student’s t-test N>15. B) Quantitative genetic interaction between genotype and histone deacetylase inhibition. The reaction norms for hypocotyl length in Col-0 (blue) and *pwr-2* (red) are shown in the presence (dashed lines, +I) or absence (solid lines) of Droxinostat. The drug x genotype interaction is shown. C) Reaction norms of the expression levels of *YUCCA8* in Col-0 and *pwr-2*. The drug x genotype interaction for Col-0 is shown. D) Hypocotyl lengths of *hda9* mutants at 22 °C or 27 °C. N>200. E) Detection of histone H3 and H2A.Z with anti-H2A.Z and anti-H3K9ac antibodies by western blots in the protein fraction that was immunoprecipitated with an anti-acetylation antibody. Detection of SUMO from the crude extract is shown as a loading control. Error bars indicate standard error. *p-values*:***<0.0001, **<0.001, *<0.05, ns=not significant.

Histone deacetylation is typically associated with down regulation of gene expression [27]. The requirement of PWR for thermomorphogenesis therefore indicates that down regulation of gene expression is also critical for proper thermal response. Thus the requirement of PWR-mediated histone deacetylation for the temperature-induced expression of *YUCCA8* and *PIF4* (Fig. 1F, Fig. 3C, S6E), suggests that these are indirect targets of the PWR/HDA9 module. To obtain further insights into *PWR-*mediated transcriptional regulation in response to temperature, we compared the *pwr-2* and Col-0 transcriptomes at 23 °C and 2-hours after a shift to 27 °C, in 6-day old seedlings. Interestingly, the number of differentially expressed genes (DEGs) between Col-0 and *pwr-2* was substantially higher at 27 °C (867 DEGs), than at 23 °C (36 DEGs) (Fig. 4A, B, Table S1 & S2). Analysis of the transcriptomes at 23 °C and 27 °C identified 30 genes to be differentially expressed in Col-0, while 623 genes to be differentially expressed in *pwr-2* (Fig. 4A). Thus, the loss of *PWR* resulted in global mis-regulation of transcription at 27 °C, which suggests that PWR dampens transcriptional response to elevated temperatures. While the majority of the mis-regulated genes were up regulated in *pwr-2* at 27 °C, consistent with the role of PWR in histone deacetylation, most of the genes that were induced by higher temperatures in Col-0 (Table S3) failed to do so in *pwr-2* mutants (Fig. S7).

**Fig. 4.**
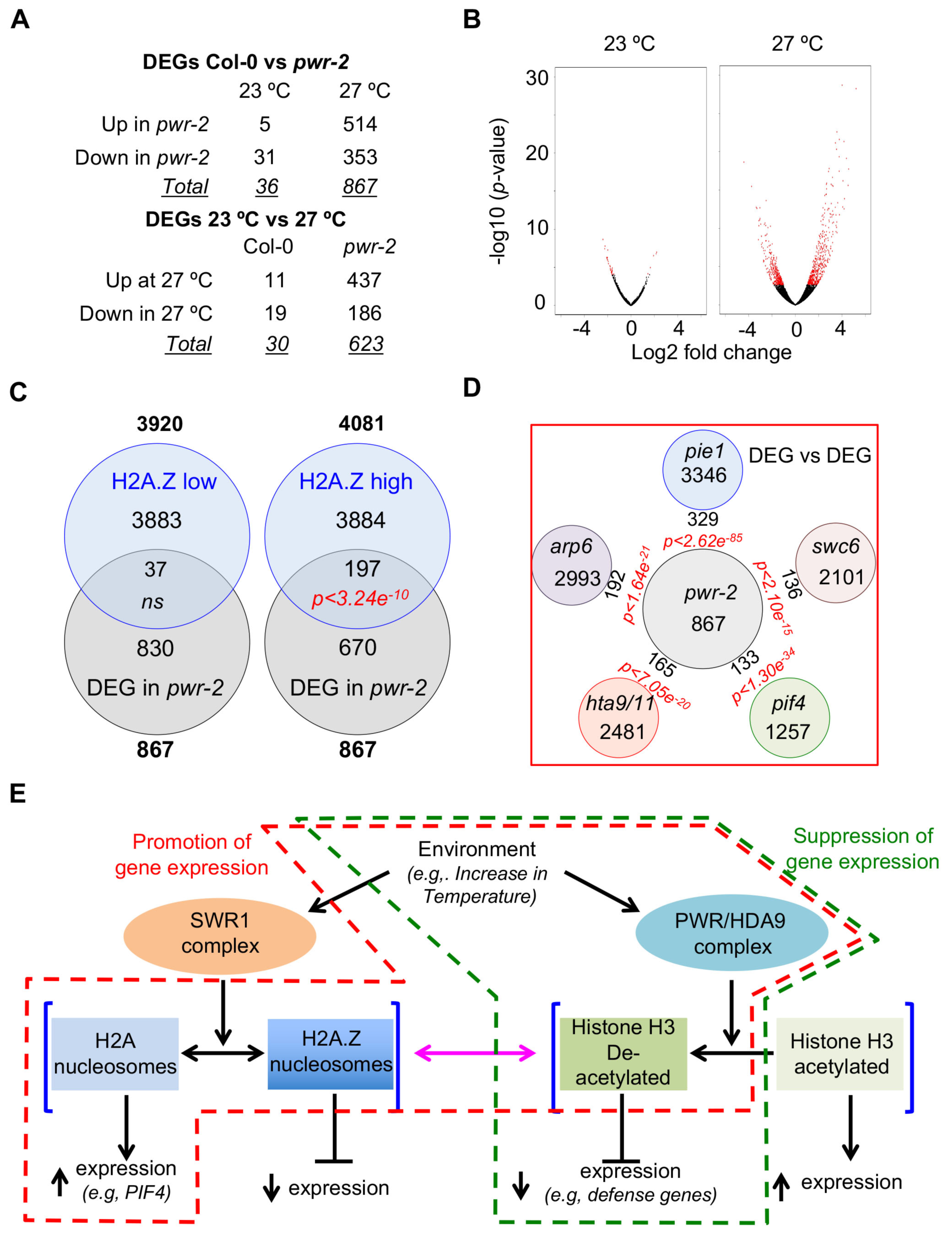
Transcriptome analysis suggests a link between *PWR* ‐mediated histone deacetylation and H2A.Z nucleosome dynamics. A) Number of DEGs (padj<0.05) between Col-0 and *pwr-2* in seedlings grown at 23°C and/or shifted to 27 °C for 2-hours. B) Volcano plots revealing massive mis-regulation of the *pwr-2* transcriptome at high temperatures. Red dots represent DEGs between *pwr-2* and Col-0. C) Overlap of DEGs between *pwr-2* and Col-0 at 27 °C with genes that are low-H2A. Z or high-H2A.Z enriched in gene bodies. Total number of DEGs is shown in bold. The H2A.Z data are from [8]. D) Overlap of DEGs between *pwr-2* and Col-0 at 27 °C with DEGs in *pie1, swc6, arp6, hta9/hta11* and *pif4.* E) A proposed model for transcriptional regulation by PWR/HDA9. The model proposes a link shown in pink between H2A.Z nucleosome dynamics and histone H3 acetylation state. The same stimulus could lead to up regulation (shown in green) or down regulation (shown in red) of gene expression, depending on the green or red paths.

Gene Ontology (GO) analysis of the DEGs between Col-0 and *pwr-2* at 27 °C showed enrichment for GO terms associated with “response” (Table S4, *Fisher Test with Yekutieli correction*). In addition to enrichment for genes associated with “response to temperature stimuli” *(p<7.4e^-6^),* various other response terms were also enriched (Table S4. e.g., response to: chemical stimuli *(p<8.7e^-59^)*, stress *(p<3.7e^-42^),* carbohydrate *(p<7.1e^-25^)*, other organism *(p<9.5e^-25^)*, water *(p<3.4e^-21^)*, defense *(p<6.3e^-21^)*, hormone *(p<1.6e^-19^)*, ethylene *(p<4.1e^-12^)),* which suggests that PWR, in addition to being involved in temperature response may be generally associated with the transcriptional regulation of “response” genes. A similar GO enrichment profile for “response” was previously reported for genes with H2A.Z enrichment in gene bodies (here after called high-H2A.Z) [8]. Therefore, we considered whether the genes that are differentially expressed in *pwr-2* overlap with H2A.Z enriched “response” genes. To assess the significance of overlaps, in addition to calculating hypergeometric probabilities, we generated 100,000 random pairs of gene lists from Arabidopsis genome, analysed the overlaps between each pairs of the gene lists, calculated their hypergeometric probability and generated a simulated distribution (Fig. S8). Through meta-analysis of the published H2A.Z data [8], with the *pwr* transcriptome data, we detected a significant overlap between DEGs between *pwr-2* and Col-0 with high-H2A.Z genes (*p<3.24e^-10^, hypergeometric probability test*), but not with low-H2A.Z genes (Fig. 4C), which suggests that the expression of a significant subset of H2A.Z enriched genes is regulated by PWR-mediated histone deacetylation. To assess whether histone acetylation is generally associated with H2A.Z enrichment, we carried out meta-analysis of publicly available data on H3K9acetylation across the genome with H2A.Z enrichment [8, 28]. While H3K9acetylation overlapped with both high-H2A.Z (850/1984, Fig. S9A, *p<2e^-7^, hypergeometric probability test*) and low-H2A.Z (1134/1984, Fig. S9A, *p<1.71e^-60^, hypergeometric probability test*) genes, we observed significant overlap of DEGs in *pwr-2* only with H3K9acetylated genes that are also enriched for H2A.Z in their gene bodies (Fig. S9B-S9D), which suggests that PWR preferentially modulates high-H2A.Z genes.

Although our findings suggested a link between PWR-mediated histone deacetylation and H2A.Z enrichment, the observed overlap with H2A.Z enriched genes could be attributed to thermal transcriptome, as temperature also affects H2A.Z nucleosome dynamics [7]. Therefore, to assess whether there is a general association between H2A.Z enrichment and PWR/HDA9 mediated transcriptional regulation, we carried out a similar meta-analysis with genes that were reported to be differentially expressed in *pwr-2* and *hda9* mutants in published studies unrelated to temperature but associated with plant aging and flowering [20, 21]. In spite of the differences in the sampled tissue, developmental states, growth conditions as well as different research groups being involved, the pattern was same, where a significant overlap was observed with high-H2A.Z but not with the low-H2A.Z genes (Fig. S9, S10A-S10C). Of the 4081 high H2A.Z genes, we detected 1068 (26%, *p<3.79e^-29^, hypergeometric probability test*) to be differentially expressed in *pwr* and/or *hda9* (S10D). These findings suggest that the association between H2A.Z enrichment and PWR/HDA9 mediated transcriptional regulation extends well beyond thermal response and hints at a general association between H2A.Z nucleosome dynamics and histone H3 deacetylation.

Histone H3 deacetylation and H2A.Z nucleosome dynamics are two fundamental, yet distinct chromatin-remodeling processes that modulate gene expression in response to diverse environmental stimuli [27, 29]. As our findings suggested a possible previously unexplored link between these two, we tested whether changes in gene expression conferred through H2A.Z nucleosome dynamics overlaps with those mediated through histone H3 deacetylation. To assess this, we compared the *pwr-2* and *hda9* transcriptomes with the published transcriptomes of H2A.Z mutants *hta9/hta11* (defective for the two out of the three H2A.Z encoding genes) [30] and the mutants for the components of the SWR1 complex *(arp6 (ACTIN RELATED PROTEIN 6), pie1 (PHOTOPERIOD INDEPENDENT EARLY FLOWERING 1) & swc6)* that mediates H2A.Z eviction/incorporation in nucleosomes. In addition, we also compared the transcriptome of *PIF4,* one of the major downstream components of the H2A.Z nucleosome dynamics [18, 31]. Here as well, despite the differences in the sampled tissue, developmental states, growth conditions and research groups involved, we observed a significant overlap between the DEGs for all the transcriptomes (Fig. 4D, Fig. S12A, S13A, S14A, S15A, S15A & S16A), which further supports the potential nexus between histone deacetylation and H2A.Z nucleosome dynamics in regulating gene expression. Analysis of up regulated and down regulated genes in each of these transcriptomes revealed a significant overlap among up-regulated genes (Fig S11AB, S12B-C, S13B-C, S15B-C & S16B-C). Nevertheless, genes whose expression changed in opposite directions also displayed an overlap (Fig S11C-D, S12D-E, S13D-E, S15D-E & S16DE), which suggests that the target genes are modulated in a dynamic manner through this interaction. Taken together, these findings further hints at a link between H2A.Z nucleosome dynamics and histone deacetylation in the regulation of gene expression in plants.

Among the analysed transcriptomes, most significant and discernible overlaps were seen with *pie1* followed by *pif4* (Fig. 4D, S11-S17), exhibiting more significance than those of *swc6, arp6* or *hta9/hta11* (Fig. 4D, S11-S17). *PIE1* also encodes a SANT domain containing protein[10], but its role in the SWR1 complex is well-studied [11, 32]. We found *pie1* mutants to display a reduced response to temperature in hypocotyl and petiole elongation, similar to *pwr-2*, although the effect was less pronounced (Fig. S18). Double mutants of *pwr-2 pie1-6* resembled the *pwr-2* mutants suggesting that *pwr* is epistatic to *pie1* (Fig. S16). Both *PIE1* and *PIF4* play critical roles in plant defense [18, 30, 33]. Defense responses are dampened at higher temperatures[4] and the up regulation of defense response genes in *pwr-2* suggests that *PWR* mediated histone deacetylation is critical in suppressing defense gene expression at elevated temperatures (Table S5).

## Discussion

We have demonstrated that *PWR* is a critical gene required for thermomorphogenesis. Previous studies suggested a key role for *PWR* in diverse developmental processes including regulation of floral determinacy, flowering and senescence [20, 21, 23]. The underlying mechanism through which PWR acts on these processes appears to be via transcriptional regulation by histone H3 deacetylation. PWR modifies acetylation status through its physical interaction with HDA9 that results in histone deacetylation at specific loci across the genome [20, 21]. Histone deacetylation has been previously shown to be essential in both developmental processes and abiotic stress response [27]. Our results demonstrate conclusively that PWR-dependent histone deacetylation is a key chromatin-remodeling mechanism required for ambient temperature-response in plants.

Chromatin remodeling through the exchange of histone H2A.Z with H2A has been previously shown to be critical in mediating thermosensory response in plants [7]. Our findings therefore reveal another layer of chromatin remodeling that is essential in mediating transcriptional responses to temperature. PWR acts at the level of chromatin in conferring thermal response and thus an upstream factor in thermosensory response in plants. Our genetic analysis supports this hypothesis. Therefore, two distinct chromatin remodeling processes *viz.,* H2A.Z nucleosome dynamics and histone H3 deacetylation appears to be essential for conferring thermal responses in plants. Nevertheless, it remains unclear as to how temperature information is perceived by the SWR1 or the PWR/HDA9 complex to regulate these chromatin-remodeling events.

Interestingly, mutations in SWR1 complex such as *arp6* and *pie1* result in contrasting temperature-induced hypocotyl phenotypes. While *arp6* appears to have longer hypocotyls even at lower temperatures [7], mutations in *pie1* result in relatively shorter hypocotyls even at elevated temperatures. Thus, the inability of *pie1* mutants to respond to elevated temperature, similar to *pwr,* reveals the complexity at the level of chromatin-remodeling governing thermal responses. It is possible that *PIE1* being a SANT-domain containing protein may have additional roles independent of the SWR1-complex. The striking overlap of *pwr-2* transcriptome with *pie1* when compared to the overlap with *arp6, swc6* and *hta9/hta11*, the distinct phenotypes of *pie1* and its genetic interaction with *pwr-2* suggest a broader role of *PIE1,* some of which are H2A.Z-independent and associated directly or indirectly with the histone deacetylation cascade regulated by PWR.

*PIE1* also plays a critical role in regulating the expression of defense genes and the role of *PIE1* in defense also differs from *arp6, swc6* and *hta9/hta11*[30, 33]. Trade-off between thermosensory growth and defense has recently been suggested to be coordinated by *PIF4*[18]. Analysis of the up-regulated genes in the *pwr-2* transcriptome also revealed an enrichment of GO terms associated with defense (Table S5). The strong overlaps of *pwr-2* transcriptome with *pie1* and *pif4* suggests that PWR-mediated histone H3 deacetylation is also critical in regulating gene expression changes that are associated with the trade off between growth and defense. Although PWR is required for PIF4 expression, we cannot rule out that PWR may also be required down stream of PIF4 in regulating defense gene expression. It is possible that PWR may act at multiple levels. It is also currently unknown whether *pwr* mutants indeed display enhanced disease resistance, which would be explored in future.

We have demonstrated that histone deacetylation is an essential aspect of thermomorphogenesis in Arabidopsis. We have also revealed that gene regulation by histone H3 deacetylation significantly overlaps with gene regulation conferred by H2A.Z nucleosome dynamics. We present a testable model that could explain our findings (Fig. 4E). We propose that histone H3 deacetylation and H2A.Z nucleosome dynamics are highly inter-connected and the response to environmental stimuli involves both processes. In addition, we hypothesise that both these processes likely influence each other. Histone H3 deacetylation regulates gene expression by altering chromatin accessibility leading to suppression of gene expression. In addition, it also affects gene regulation by potentially modulating H2A.Z eviction/incorporation dynamics. Thus, both up/down regulation of gene expression could result from same stimuli due to intra-nucleosomal interactions at the chromatin. Similar suggestions have been made in other systems. It has been suggested that histone H3 acetylation patterns can modulate H2A.Z nucleosome dynamics in yeast [34], whereas H2A.Z has been shown to promote H3 and H4 acetylation in mammalian cells [35, 36]. Exploring the mechanistic basis of this connection would be an exciting avenue for future work.

## Materials and Methods

### Plant material and phenotyping

All mutants were in the Col-0 background unless otherwise specified. All T-DNA insertion lines as well as most of the mutant lines used in this study were obtained from the European Arabidopsis Stock Centre. *pwr-1*, *pwr-2, hda9-1, hda9-2, hda6* and *hda19* mutants have been described [23, 37-39]. *pwr-1* and *pPWR::PWR-GFP* lines were gifted by Prof. Xuemei Chen and *pif4-101* is from Prof. Christian Fankhauser. All double mutants were obtained by crossing and confirmed by genotyping. Hypocotyl and petiole length measurements were done as described previously[40]. Briefly, seeds were sterilised, sown on Murashige-Skoog media and then stratified for 2 days at 4°C in dark. The plates were then transferred to CU41L5-Percival growth chambers (Percival Inc, Canada) at 23° C or 27° C in short day conditions (8 hour light / 16 hour dark) and grown vertically for 10 days. For the T-DNA screening, more than 20 seedlings representing each of the 5000 T-DNA lines were grown at 23 °C and 27 °C and the seedlings were visibly inspected for attenuated response. To quantify the hypocotyl elongation, subsequently, plates with plantlets were imaged and the hypocotyl length was measured using Image J (NIH). All T-DNA lines used in subsequent analysis described in this study were confirmed by using T-DNA insertion using primers listed in the Table S6. Flowering time measurements were done as described previously and total leaf number is used as a proxy for flowering time [3]. For the HDAC inhibitor assays all the compounds (Sigma-Aldrich) were dissolved in the described concentration in DMSO and the solvent lacking the compounds was used as a mock control.

### DNA/RNA Analyses

DNA and RNA extractions were done as described previously [41]. For gene expression studies DNAse I (Roche)-treated 1ug of total RNA was used for cDNA synthesis using the First strand cDNA synthesis kit (Roche) and the resulting cDNA was diluted and used for realtime PCR analysis with a Lightcycler 480 system (Roche) with SYBR green. The specific primers used for real-time PCR analysis are in Table S6. Relative expression levels were obtained using the ΔΔcT method [42] using either *UBIQUITIN* or *TUBULIN* as internal controls.

### Immunoprecipitation

Approximately fifty 3-week old *A.thaliana* seedlings were ground in liquid nitrogen and 1mL of incubation Buffer (50mM Tris-Hcl pH7.5, 150mM NaCl, 10mM EDTA, 0.2% Triton, Roche protease inhibitor cocktail) was added. The samples were spun for 15 min at 4 °C at 13200 rpm and 50μL of supernatant kept for crude extract analysis. 250ng of anti-acetylated Lysine antibody (mouse anti-acetylated-Lysine antibody Ac-K-103, Cell Signaling Technology; dilution 12000) were added to each sample and incubated with continuous shaking for 2h at 4 °C. 50 μL of protein A agarose beads were then added for 2 more hours of incubation in the same conditions. The resin was washed 3x with 1mL of incubation buffer and denaturated for 3min at 95 °C in presence of 50 μL of Laemmli buffer [43] before immunodetection through western blots.

### Western blots

Western blots were done as described previously [44]. Equal amounts of protein were loaded onto a sodium dodecyl sulfate-polyacrylamide gel electrophoresis (SDS-PAGE), followed by electrophoresis and transfer to Protran BA85 nitrocellulose membranes (Whatman, Germany). Transferred proteins were visualized by Ponceau S red staining. Plant protein samples obtained from *A. thaliana* (20 seedlings), were homogenized in 250 μL of Laemmli loading buffer [43]. Antibodies used for Western blotting were anti-H2A.Z (dilution 1/5000), anti-SUMO1 (dilution 1/5000), anti-H3K9ac (dilution 1/5000) goat anti-mouse and anti-rabbit IgG-HRP antibodies (Santa Cruz, dilution 110000).

### Transcriptome studies

RNA-seq analysis was done as described previously [45]. About one hundred 6-day-old seedlings of Col-0 and *pwr*-2 each were grown at 23 °C in short days (SD) in growth chambers (GR-36, Percival Scientific, Canada). Half of the samples were moved to 27 °C. Tissue from whole seedlings were collected for RNA extraction from both 23 °C and 27 °C after 2-hours. Two biological replicates were used. Total RNA was extracted from two biological replicates using Isolate II RNA plant kit (Bioline Pty Ltd, Australia). The libraries were prepared on an Illumina HiSeq^TM^ 2000 platform using paired-end sequencing of 90 bp in length at BGI-Shenzen (Beijing Genomics Institute). *FastQC* (http://www.bioinformatics.babraham.ac.uk/projects/fastqc) was used to perform the initial quality control check of the transcriptome data. *SortmeRNA* was used to filter the rRNA sequences from the datasets, using its default rRNA databases comprising of 16S, 18S 23S and 28S rRNAs[46]. The reads for each sample were aligned to *Arabidopsis thaliana* TAIR 10 genome using *Tophat2 (v2.1.0)* [47] and *bowtie2 (v2.1.0)* [48]. Raw abundance counts were obtained from the Bio conductor-R-subread package using *featureCounts (v1.4.5)* [49] from the output produced by *Tophat2.* Only fragments with both reads successfully aligned (specified through ‐p and ‐B parameters in *featureCounts*) were considered for summarization. The resulting lists of abundance counts were used as an input data for *DESeq2 (v1.14.1)* [50] differential expression analysis pipeline. For differential expression analysis and estimation of dispersions across libraries in *DESeq2*, batch effect between replicates was accounted for through a negative binomial GLM as described previously [45]. Genes with a padj<0.05 (Benjamini-Hochberg corrected p-values) were termed as differentially expressed genes (DEGs). The gene lists generated through the analysis of differential expression were used in the online program AgriGO to identify enriched GO terms [51]. Additional gene lists for overlap analysis were either obtained from published data [8, 18, 20, 30, 33]. Overlaps between gene lists were tested through hypergeometric probabilities as well as through simulation studies performed in R. 100,000 random gene lists of comparable sizes to the gene lists that were analysed (500-2000 genes) were generated from Arabidopsis. The hypergeometric probabilities for the overlaps observed in the simulated dataset were calculated using hypergeometric probability function in R.

## Acknowledgments

We thank the European and North American Arabidopsis stock Centres, and Xuemei Chen for the seeds. We thank Iain Searle, John Alvarez, David Smyth and members of the SKB lab for critical comments on the manuscript. The RNA-seq data presented in this paper is available in GEO repository with the accession number GSE101782.

## Author contributions

CT, MvZ and SB designed and conceived the study. CT, SS, RS, LvdW and MN performed experiments. CT, ASY, DT and SB analysed the data. CT, ASY and SB wrote the manuscript with inputs from all authors.

## Supporting Information

**Fig. S1.**
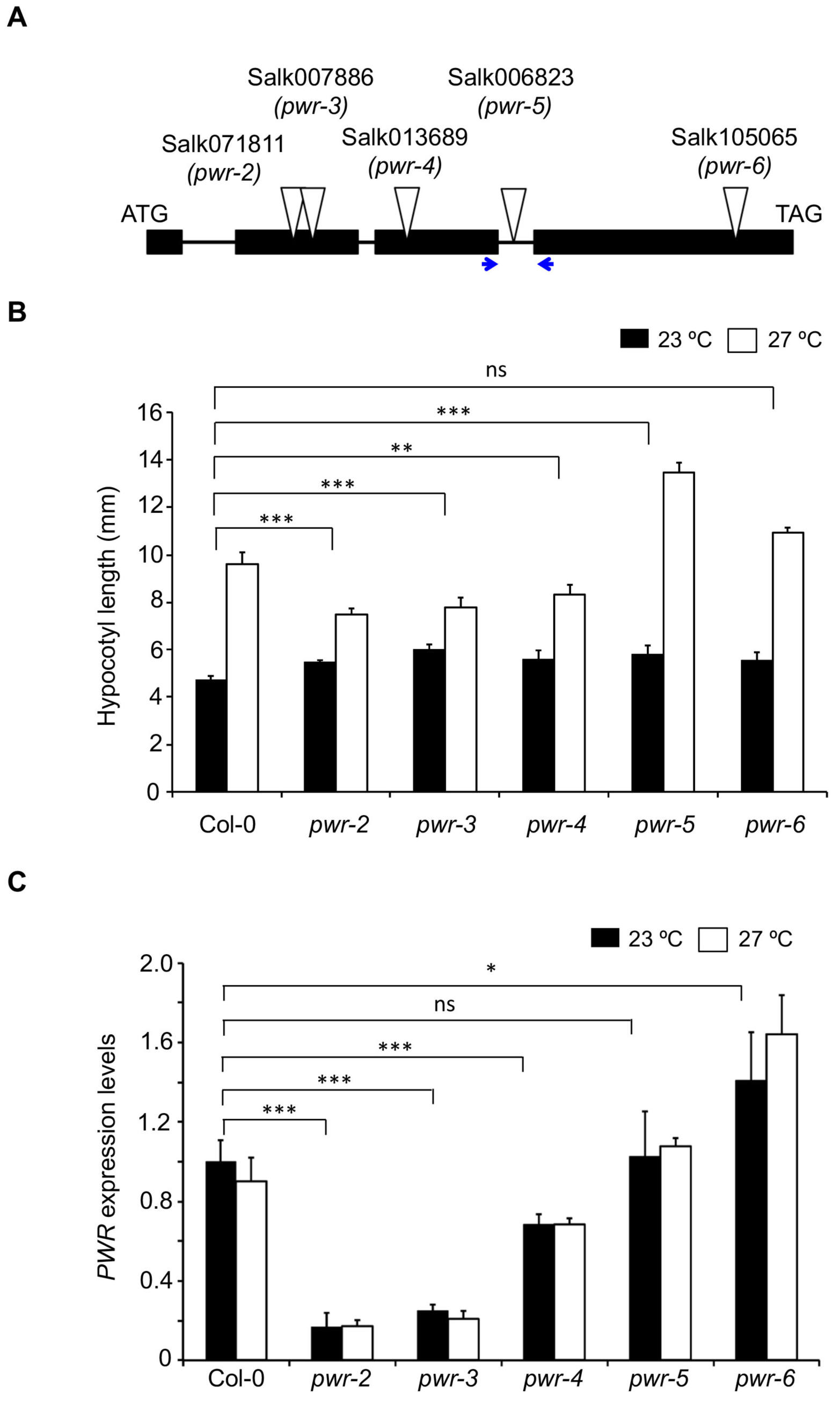
Identification of other *pwr* alleles. A) T-DNA insertion lines at the *PWR* locus in the Col-0 background, the location of the insert and their corresponding Salk identifiers. Of these *pwr-2* has been previously described [23]. The location of primers used to analyse expression is shown in blue. B) Hypocotyl lengths of different *pwr* alleles at 23 °C and 27 °C short days. The *p-values* for G x E between the different *pwr* alleles and Col-0 is shown. C) Relative expression levels of *PWR* different mutant alleles grown 23 °C and 27 °C short days. The *pwr-5* and *pwr-6* alleles have higher *PWR* expression and thus are not RNA-null alleles. The *p-values* for the difference in *PWR* expression between *pwr* alleles and Col-0 at 23 °C determined through a Student’s t-test is shown. No significant differences were observed between 23 °C and 27 °C. Error bars indicate standard error. *p‐ values*: ***<0.0001, **<0.001, *<0.05, ns=not significant.

**Fig. S2.**
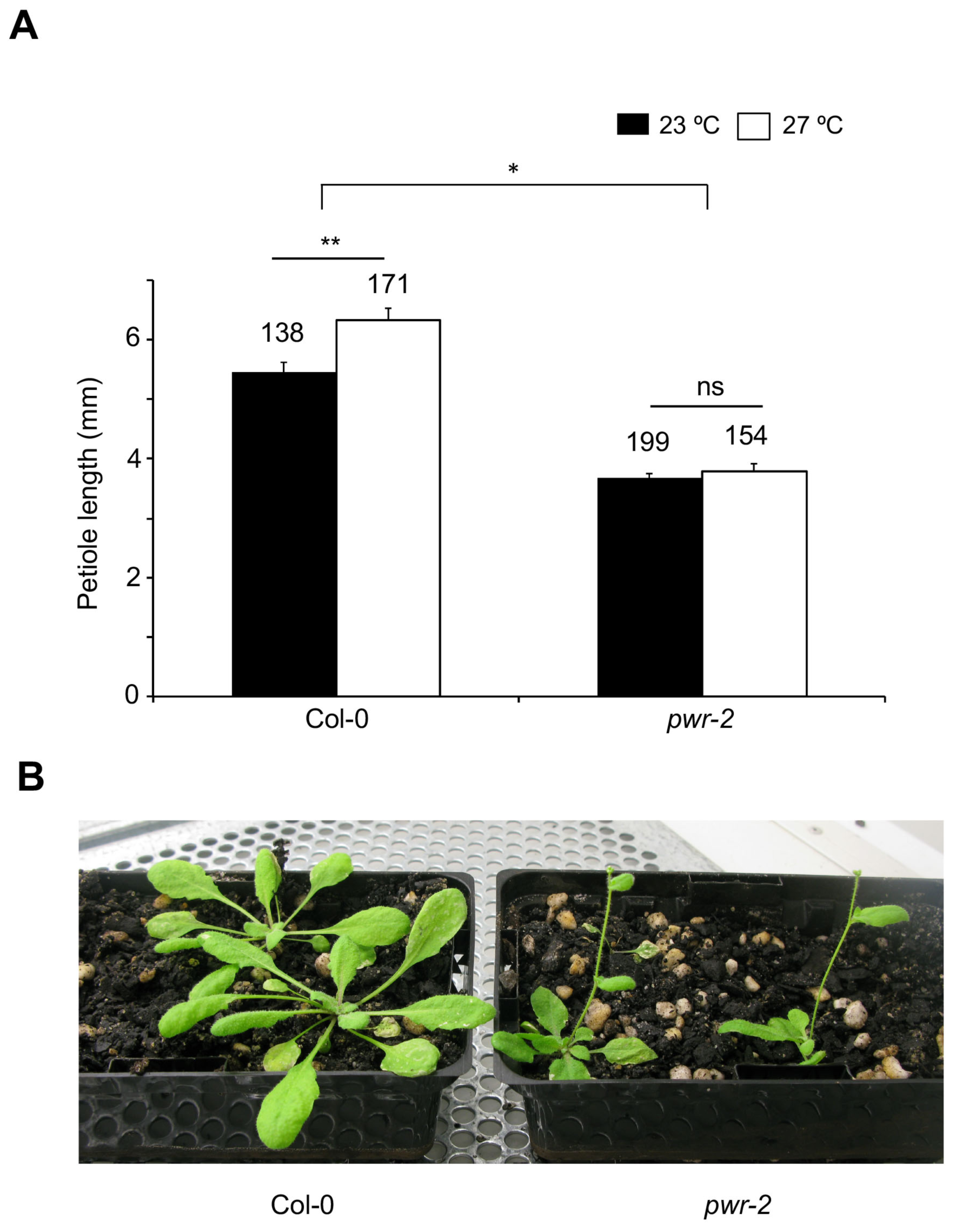
Growth differences between Col-0 and *pwr-2* at 27 °C. A) Petiole length of Col-0 and *pwr-2* at 23 °C and 27 °C. The number of petioles measured is shown above the bars. The *p‐ values* for the difference in petiole lengths between temperature determined through Student’s ttest is shown above the bars. The *p-value* for G x E interaction is also shown at the top. B) *4-*week old Col-0 and *pwr-2* plants grown at 27 °C in short days. Note the compact stature and early flowering in *pwr-2* compared to Col-0. Error bars indicate standard error. *p‐ values*: ***<0.0001, **<0.001, *<0.05, ns=not significant.

**Fig. S3.**
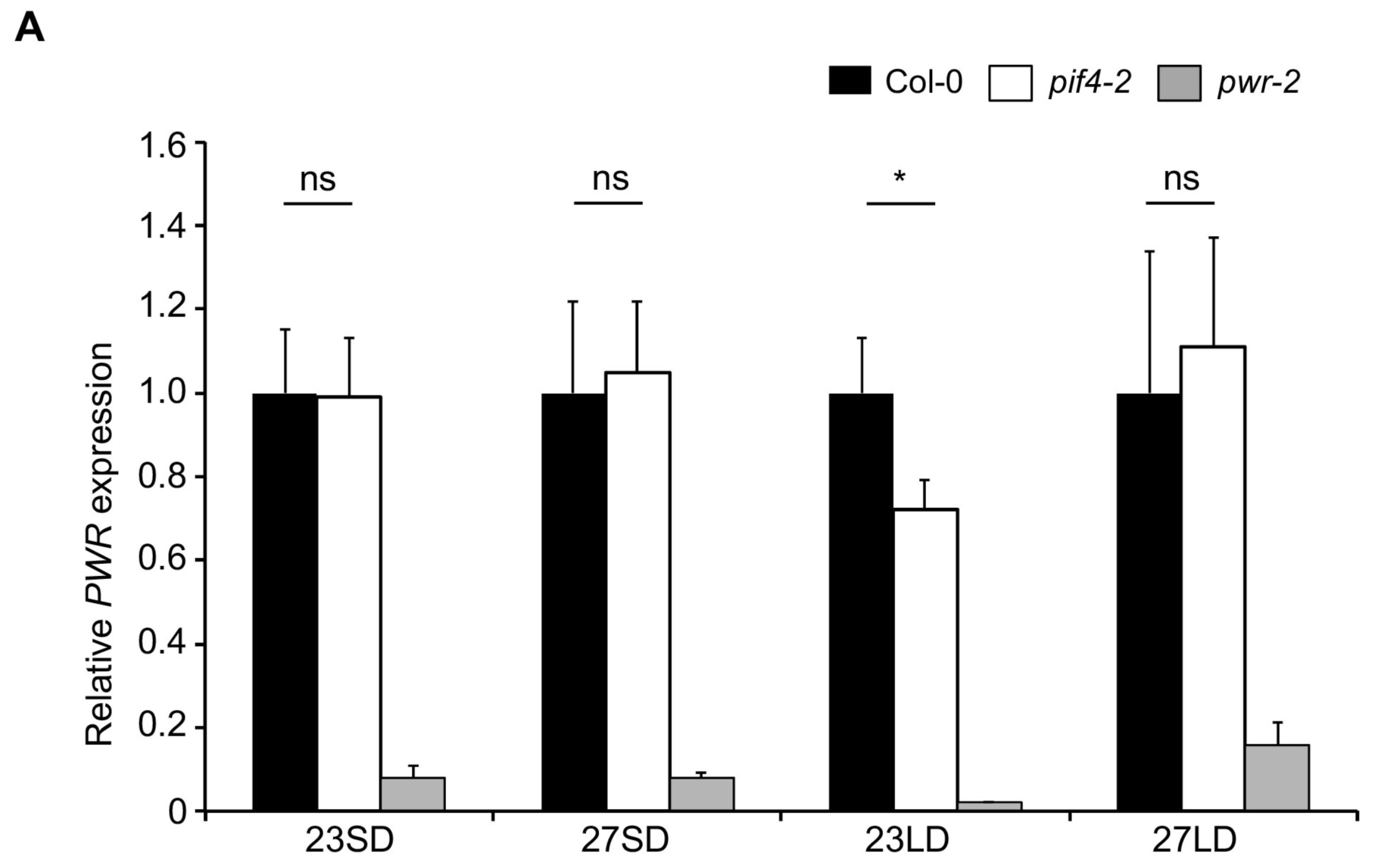
*PWR* expression is unaffected in *pif4-2* mutants. Relative expression levels of *PWR* in Col-0 (black) and *pif4-2* (white) at different temperatures (23 °C and 27 °C) in long (LD) and short (SD) days are shown. Expression is normalized against the *PWR* expression levels in Col-0 for each condition and *TUBULIN* was used as an internal control. *PWR* expression levels in *pwr-2* (grey) are shown as a negative control. Error bars indicate standard error. *p‐ values*: ***<0.0001, **<0.001, *<0.05, ns=not significant.

**Fig. S4.**
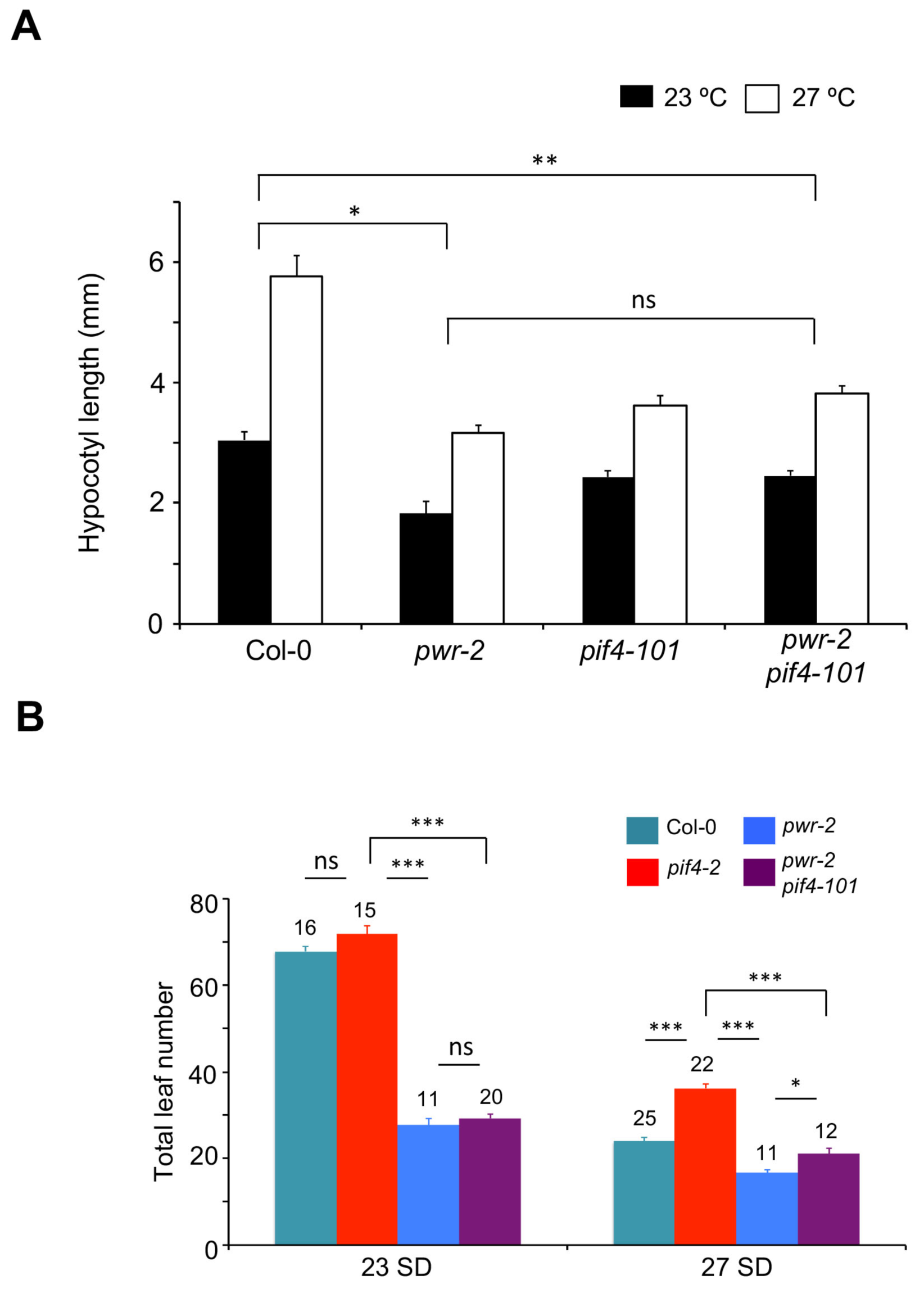
Analysis of the genetic interactions between *pwr-2* and *pif4-101.* A) Hypocotyl lengths of various genotypes at 23 °C and 27 °C*. p-values* for the corresponding GxE interactions determined through ANOVA are shown. B) Flowering time of *pif4-2 pwr-101* double mutants compared to single mutants at two different temperatures. Number of plants and the *P-values* determined through Student’s t-test are shown above bar graphs. Error bars indicate standard error. *p‐ values*: ***<0.0001, **<0.001, *<0.05, ns=not significant.

**Fig. S5.**
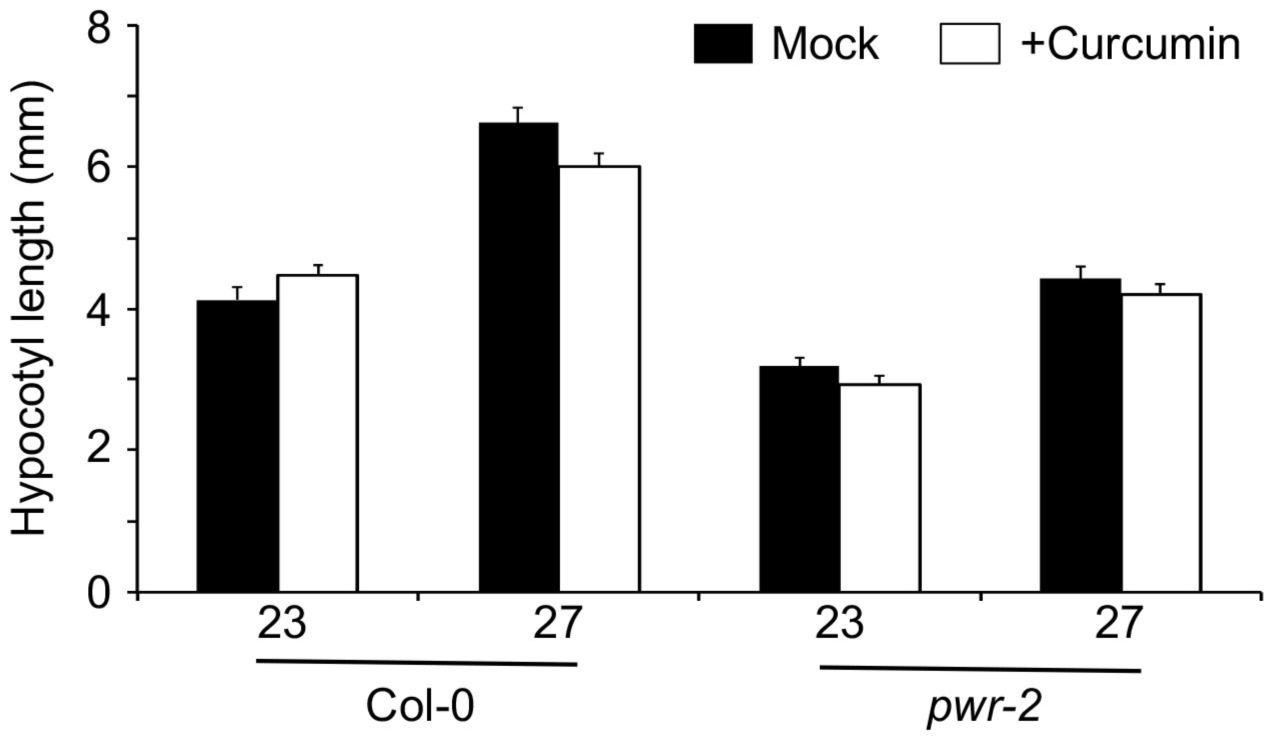
Inhibition of histone acetylation does not affect thermomorphogenesis. Hypocotyl elongation in Col-0 and *pwr-2* mutants observed in plants grown in presence of 10uM of curcumin, a inhibitor or histone acetyl transferase compared to mock at 23 °C and 27 °C.

**Fig. S6.**
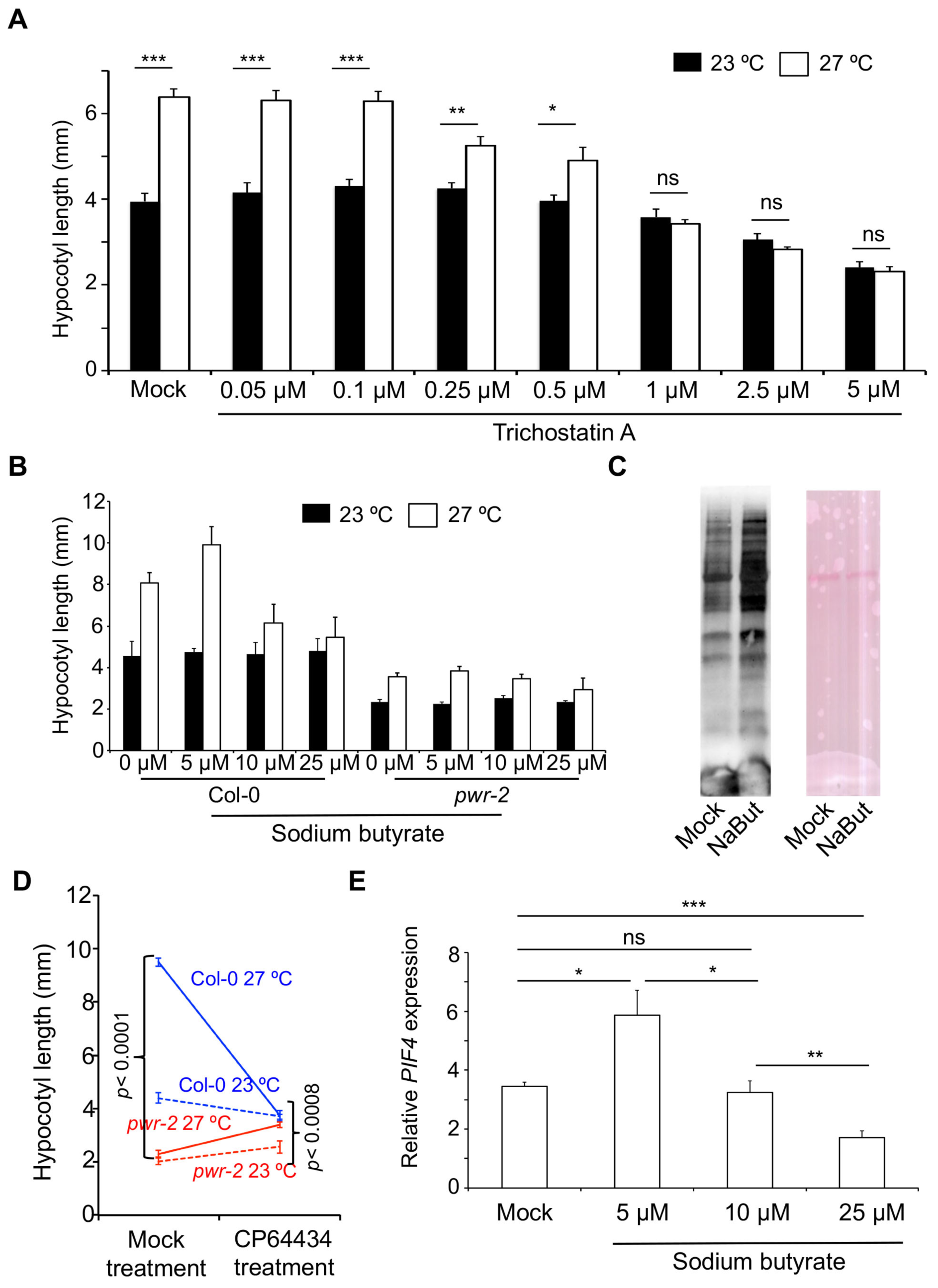
Histone deacetylation is essential for thermomorphogenesis. A & B) Dose-response effect of Trichostatin-A (A, N=9 replicates with >20 plants), and Sodium Butyrate (B, N=8), on Col-0 (A, B) and *pwr-2* (B) at 23 °C and 27 °C. *p-values* shown are derived from one way ANOVA using the presence/absence of the drug as a factor. C) Western blots of crude plant extracts probed with anti-acetylated antibody from plants grown with or without Sodium Butyrate. Ponceau-S stained gel is shown as loading control. D) Reaction norms of hypocotyl lengths in Mock vs HDAC inhibitor treatment. The *p-value* for the drug x genotype interaction, determined through ANOVA is shown. E) Effect of inhibition of histone deacetylation on *PIF4* expression in seedlings grown at 23 °C or 27 °C with or without Sodium Butyrate at different concentrations. *p-values* are determined by one way ANOVA with temperature as a factor. Error bars represent standard error. *p‐ values*: ***<0.0001, **<0.001, *<0.05, ns=not significant.

**Fig. S7.**
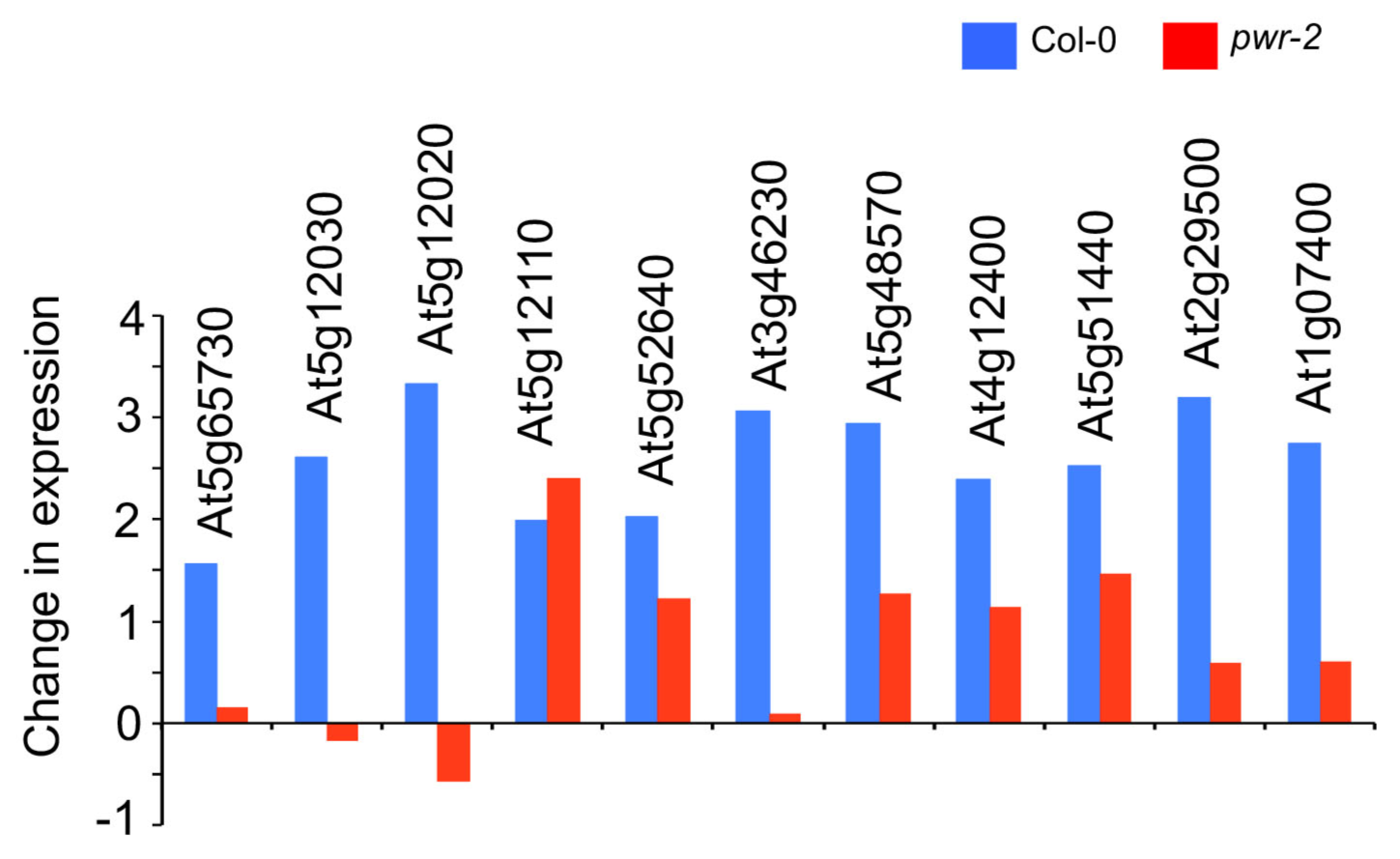
Genes up regulated in Col-0 at higher temperature are mostly attenuated in *pwr-2.* Comparison of the fold change in expression levels for the 11 genes that are up regulated at 27 °C in Col-0 (blue) and their corresponding response in *pwr-2* (red).

**Fig. S8.**
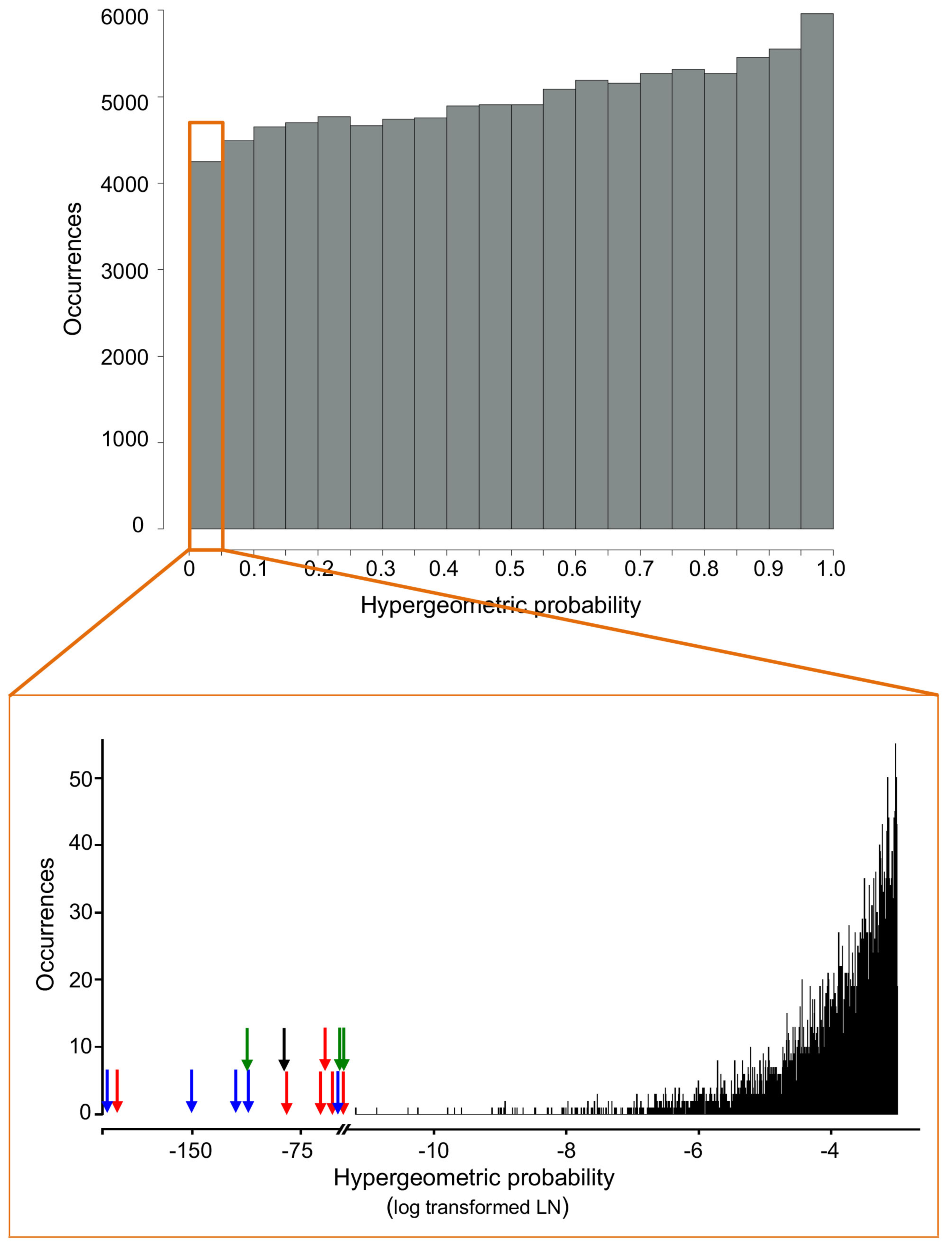
Distribution of hypergeometric probabilities obtained from 100,000 simulations of gene overlap analysis by sampling Arabidopsis genome. The top panel represents the frequency distribution of hypergeometric probabalities obtained through simulations. Random gene lists of 500 to 2000 genes were generated by sampling the Arabidopsis genome and the hypergeometric probability was estimated for the gene overlap. The entire analysis is repeated 100,000 times and the distribution of the probabilities is shown. To demonstrate the significance of the overlaps that are shown in this paper, the p-value distribution for those tests that yielded p<0.05 is shown in the second panel (expanded orange box). The p-values have been log-transformed to show the magnitude of the differences. Please note the differences in the scale for X-axis. The p-values for the gene overlaps for *pwr-2* and *hda9* from different labs is shown by green and black arrows respectively. The red-arrows depict the p-values for the gene overlap with DEGs in *pwr-2* from our group and the blue arrows refer to the same for the overlap between the combined dataset for *pwr-2* and other genes. Note all the p-values fall completely outside the distributions obtained through simulations.

**Fig. S9.**
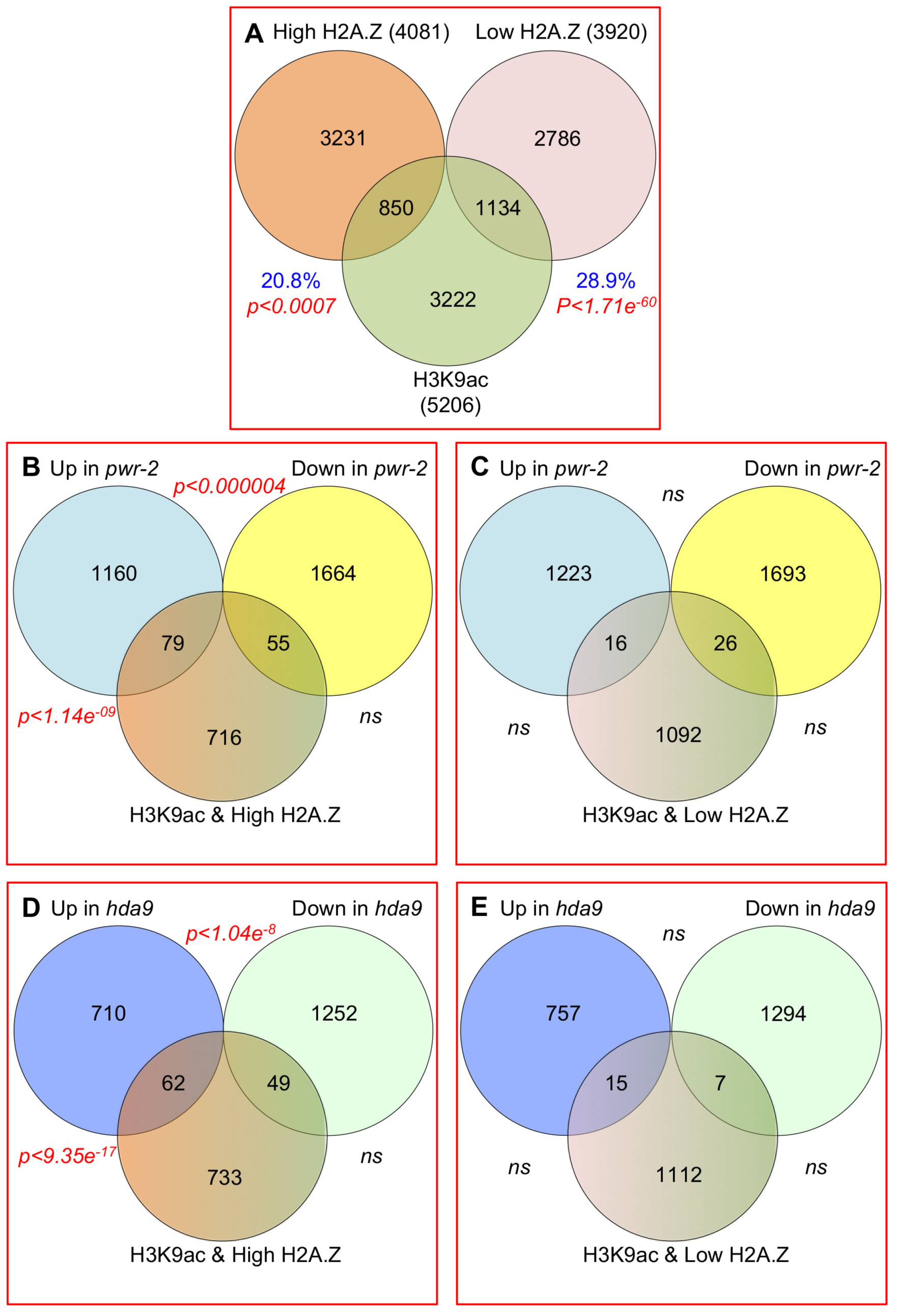
Overlap between H2A.Z enrichment, H3K9acetylation and PWR-dependent gene regulation. A) Overlap between genes with H3K9 acetylation and genes with either high or low H2A.Z in their gene bodies B & C) Overlap among genes that are up/down regulated in *pwr-2* with H3K9acetylated genes with high (B) or low (C) H2A.Z. The DEGs data is the union of Tasset et al (current study) and Kim et al [21] study from seedlings. D & E) Overlap among genes that are up/down regulated in *hda9* with H3K9acetylated genes with high (B) or low (C) H2A.Z. The DEGs data is from Kim et al [21]. The significant p-values shown in red represent hypergeometric probability for the overlap. ns = not significant.

**Fig. S10.**
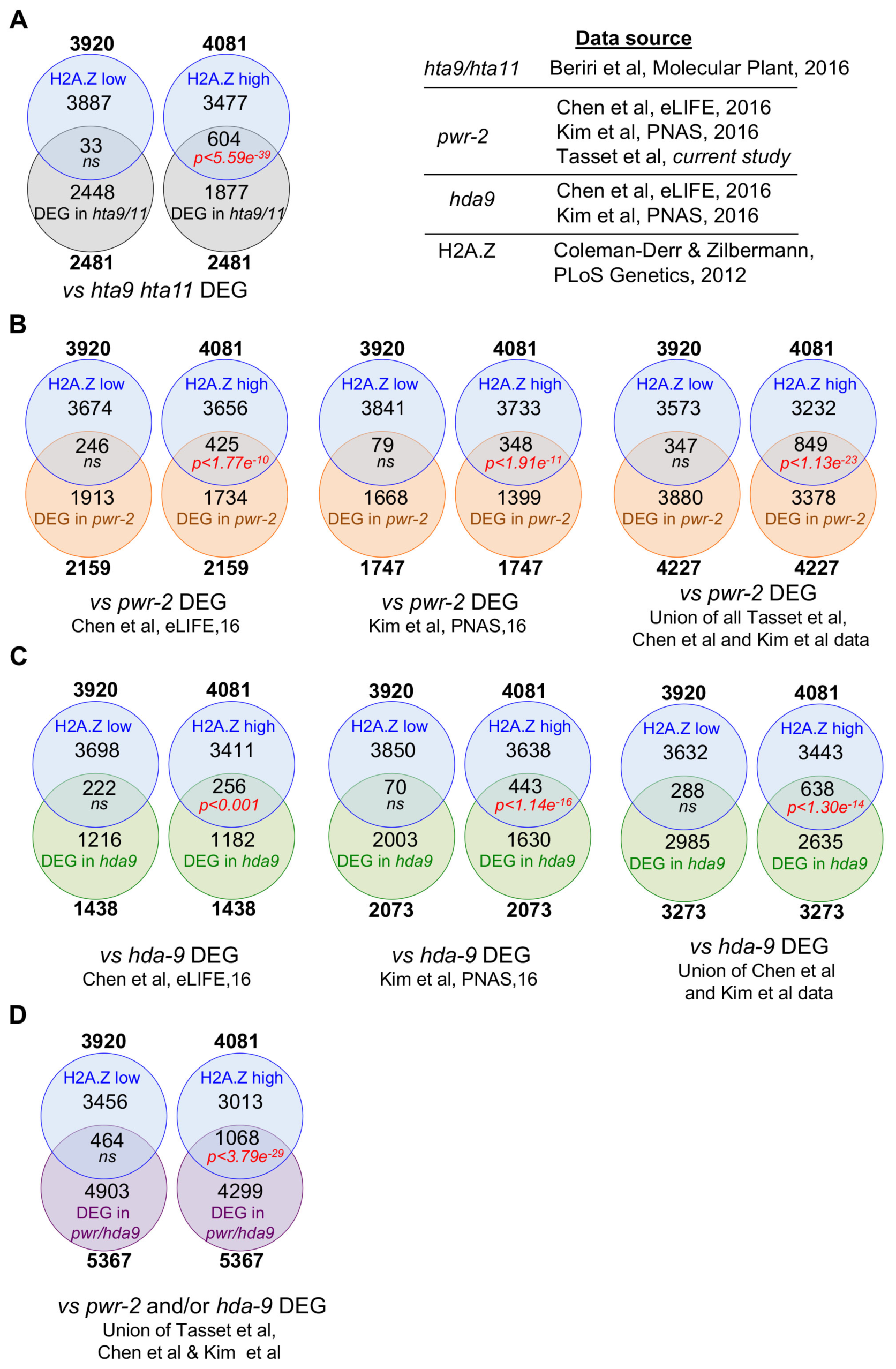
Overlap analysis of DEGs with genes that are low-H2A. Z and high-H2A.Z genes. A) Overlap of the DEGs in *hta9/hta11* double mutants with low-H2A.Z and high-H2A.Z genes. B) Overlap of DEGs in *pwr-2* mutants from two different data sets with low-H2A.Z and high-H2A.Z genes are shown. C) Overlap of DEGs in *hda9-1* mutants from two different data sets with low-H2A.Z and high H2A.Z genes are shown. D) Overlap of DEGs in *pwr-2* and/or *hda9* with low-H2A.Z and high-H2A.Z genes are shown. Total numbers of genes in respective gene lists are shown in bold. The data source is shown on top. *p-values* refer to hypergeometric probabilities and the significant *p-values* are shown in red. ns = not significant.

**Fig. S11.**
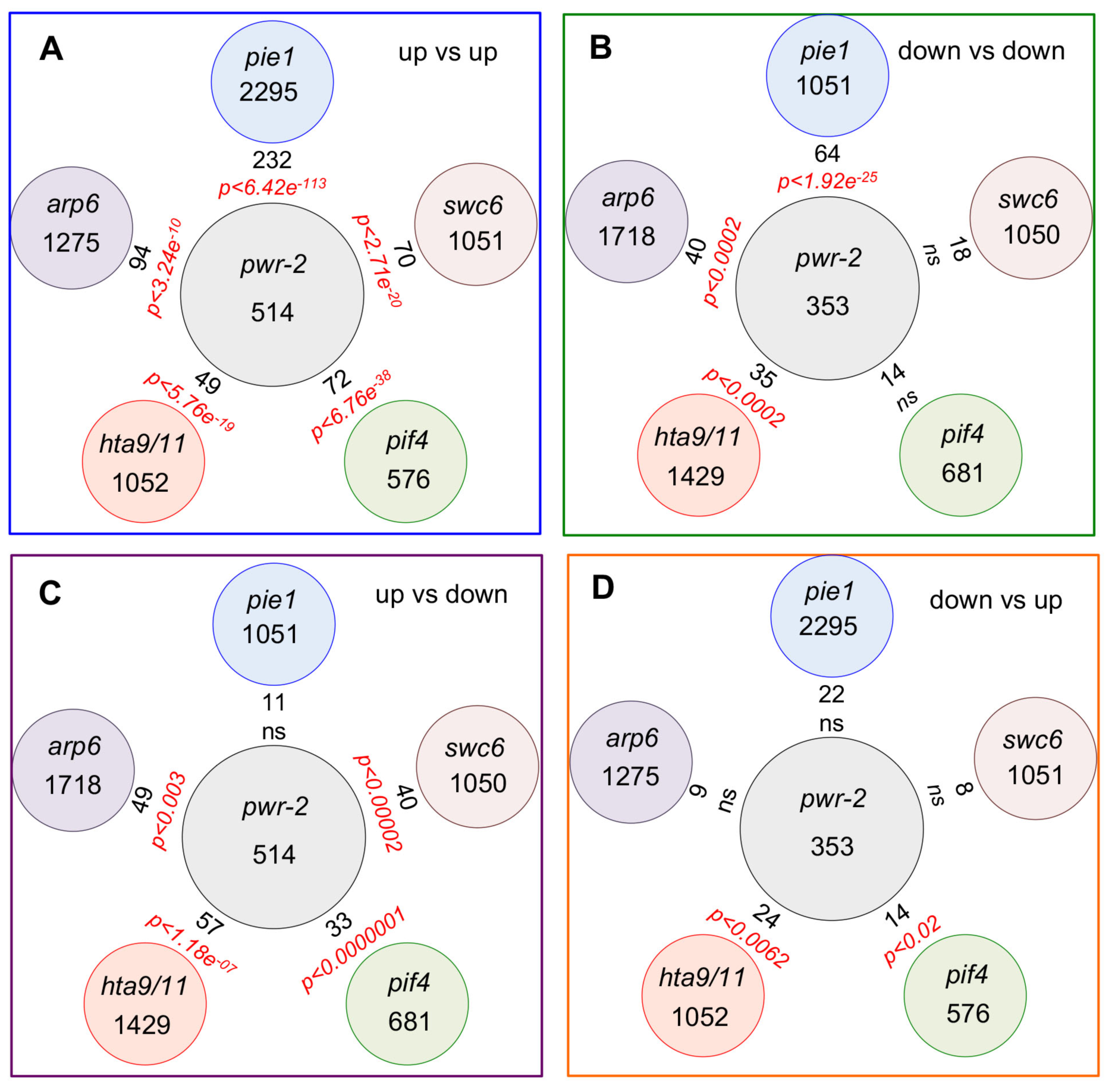
H2A.Z-related transcriptional response overlaps with that of *PWR.* A-E) Overlap of DEGs in *pwr-2* compared to Col-0 at 27 °C with DEGs in *pie1, swc6, pif4, hta9/hta11* and *arp6.* A) Overlap among DEGs. B) Overlap among genes that are up regulated in all genotypes. C) Overlap among genes that are down regulated in all genotypes. D) Overlap among genes that were up regulated in *pwr-2,* but down-regulated in other genotypes. E) Overlap among genes that were down-regulated in *pwr-2*, but up regulated in other genotypes. The total number of DEGs is shown in circles and the numbers in between represent the overlapping set of genes. The significant p-values shown in red represent hypergeometric probability for the overlap. ns = not significant. The transcriptome data is from[18, 30, 33]

**Fig. S12.**
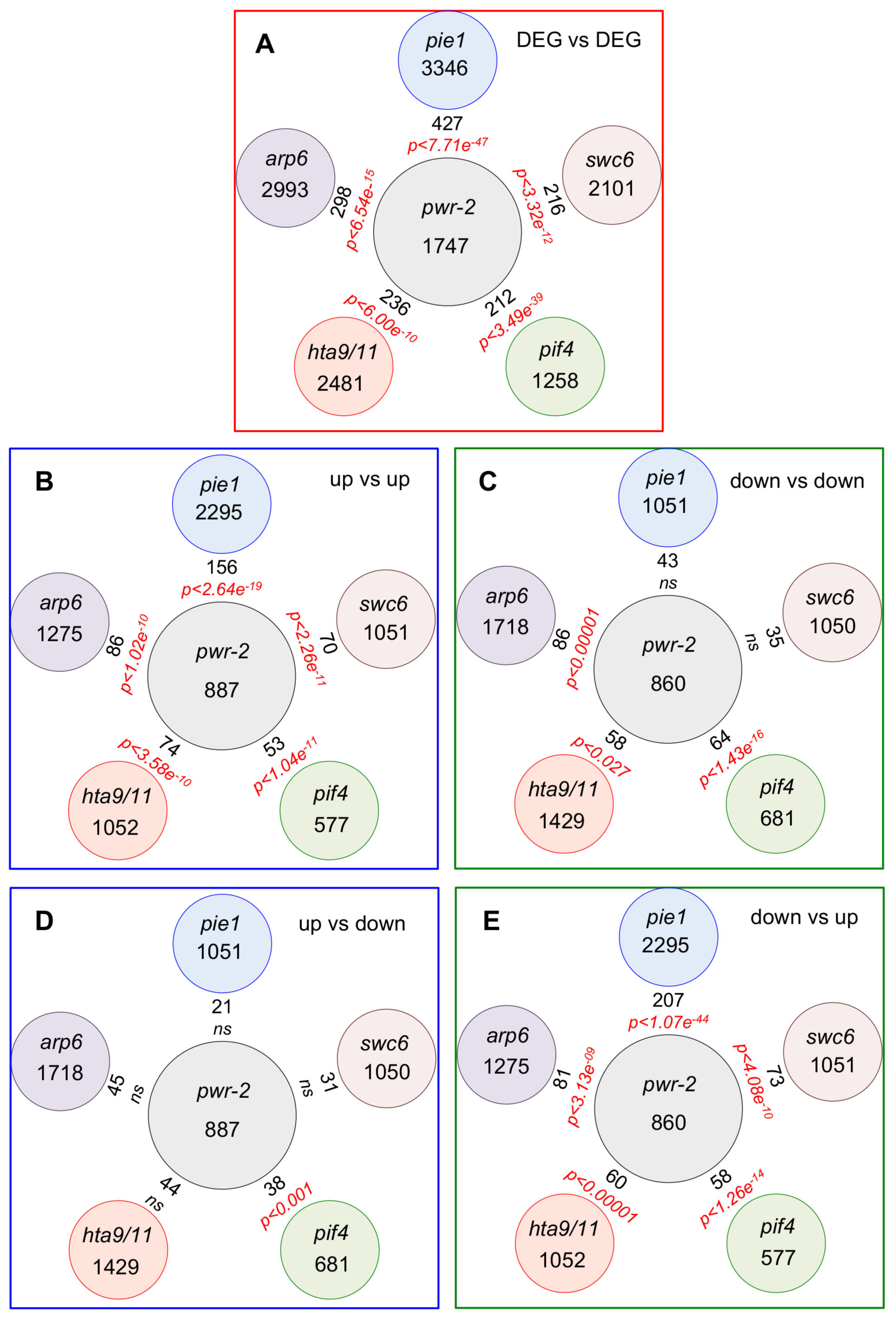
Overlap analysis of DEGs in *pwr-2* with DEGs in the transcriptomes of *pie1, swc6, arp6, pif4-2* and the *hta9/hta11* double mutants based on Chen et al data. A-E) Overlap of DEGs in *pwr-2* compared to Col-0 with DEGs in *pie1, swc6, pif4, hta9/hta11* and *arp6.* A) Overlap among DEGs. B) Overlap among genes that are up regulated in all genotypes. C) Overlap among genes that are down regulated in all genotypes. D) Overlap among genes that were up regulated in *pwr-2,* but down regulated in other genotypes. E) Overlap among genes that were down regulated in *pwr-2*, but up regulated in other genotypes. The total number of DEGs is shown in circles and the numbers in between represent the overlapping set of genes. The significant p-values shown in red represent hypergeometric probability for the overlap. ns = not significant. The transcriptome data is from [18, 30, 33]

**Fig. S13.**
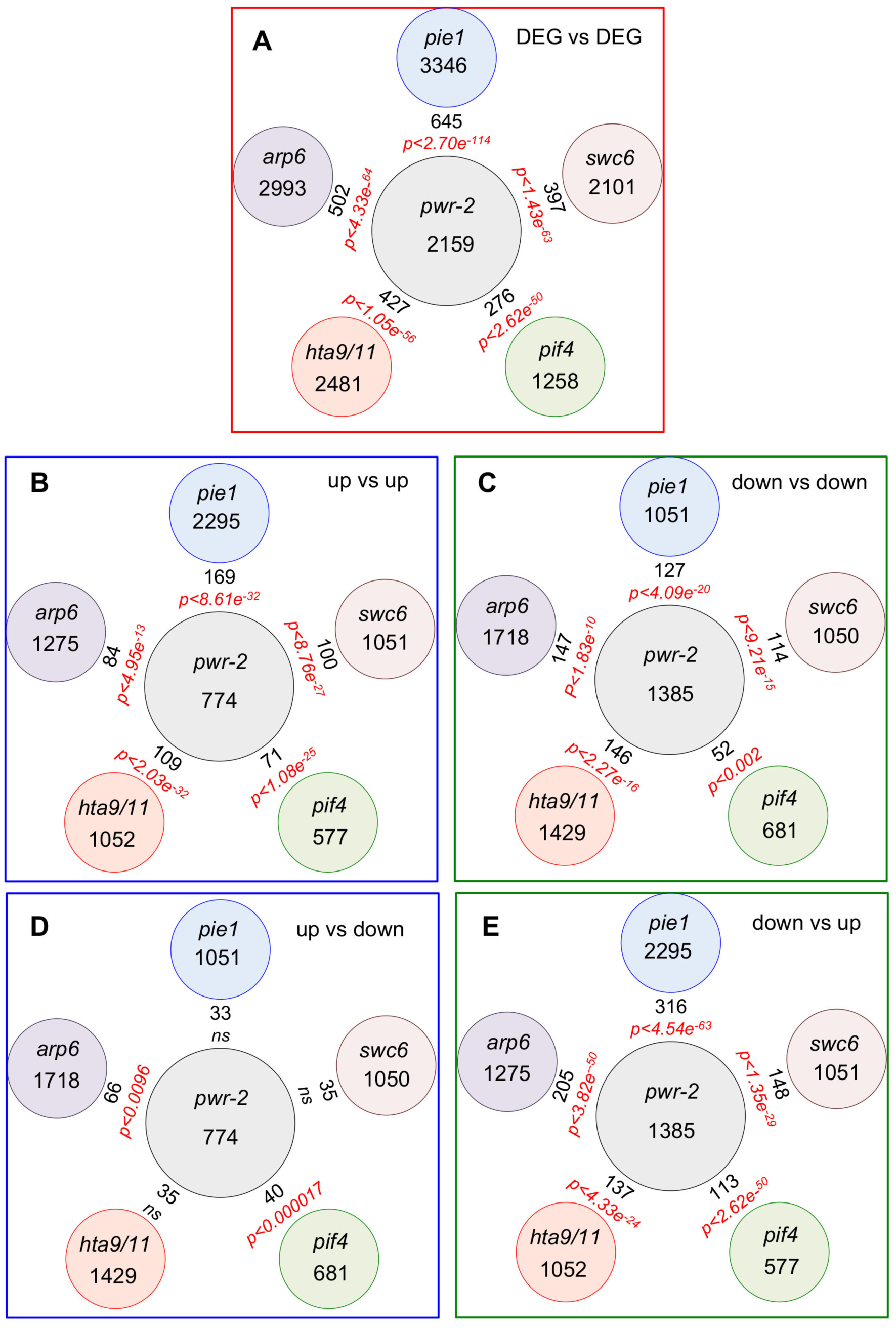
Overlap analysis of DEGs in *pwr-2* with DEGs in the transcriptomes of *pie1, swc6, arp6, pif4-2* and the *hta9/hta11* double mutants based on Kim et al data. A-E) Overlap of DEGs in *pwr-2* compared to Col-0 with DEGs in *pie1, swc6, pif4, hta9/hta11* and *arp6.* A) Overlap among DEGs. B) Overlap among genes that are up regulated in all genotypes. C) Overlap among genes that are down regulated in all genotypes. D) Overlap among genes that were up regulated in *pwr-2,* but down regulated in other genotypes. E) Overlap among genes that were down regulated in *pwr-2*, but up regulated in other genotypes. The total number of DEGs is shown in circles and the numbers in between represent the overlapping set of genes. The significant p-values shown in red represent hypergeometric probability for the overlap. ns = not significant The transcriptome data is from [18, 30, 33]

**Fig. S14.**
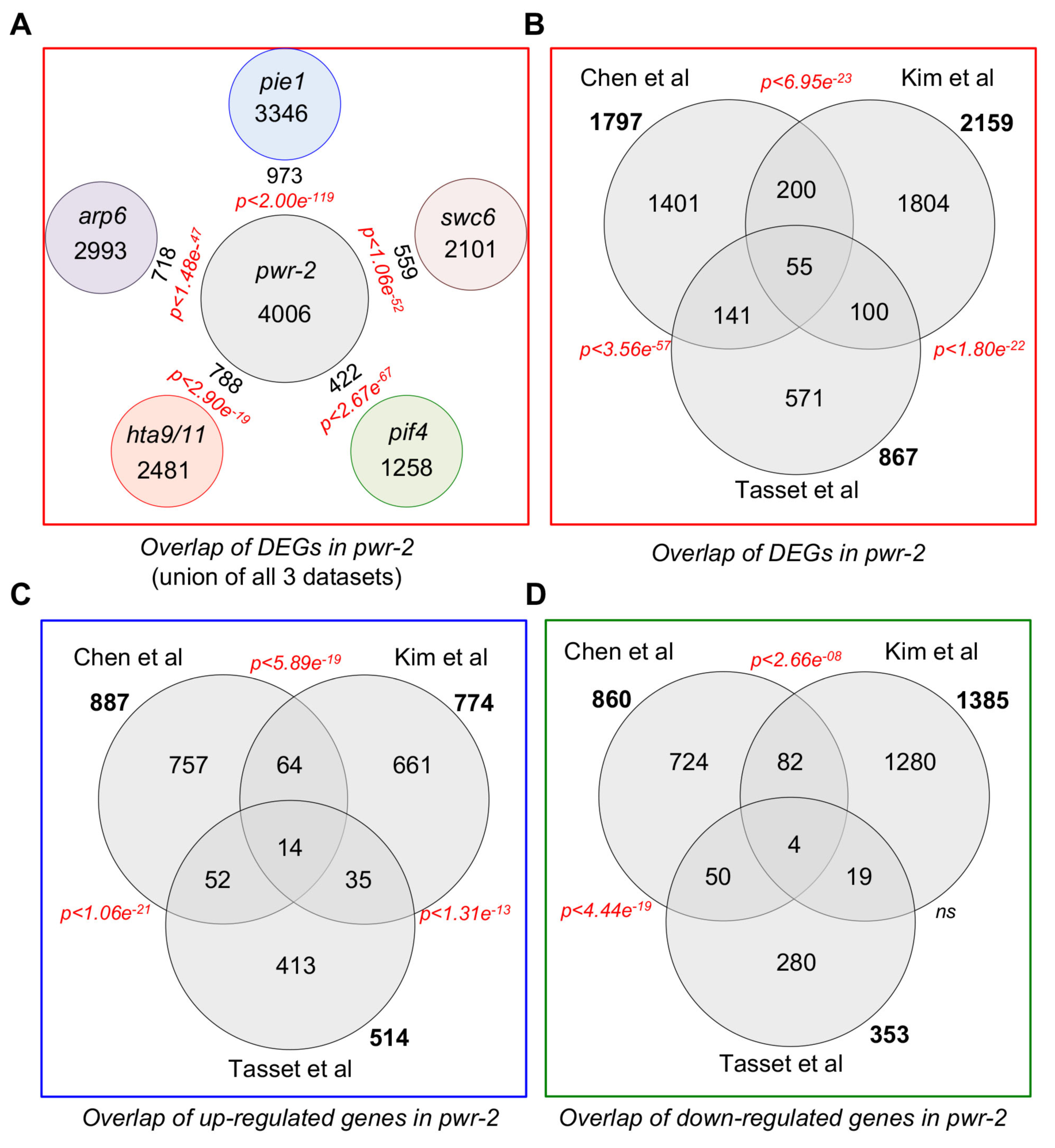
Comparison of the overlaps between the transcriptional response observed in *pwr-2* in three different datasets and their overlap with the DEGs in *arp6, pie1, swc6, hta9/hta11* and *pif4.* A) Overlap of DEGs in *pwr* compared to Col-0 compiled from all three datasets (excluding genes that did not change in the same direction in the datasets) with DEGs in *pie1, swc6, pif4, hta9/hta11* and *arp6.* The total number of DEGs is shown in circles and the numbers in between represent the overlapping set of genes. The transcriptome data is from [18, 30, 33]. B) Overlap of the DEGs in *pwr* in the three different datasets. The transcriptome data is from [20, 21] The p-values are shown next to each of the overlaps. C) Overlap among up-regulated genes. D) Overlap among down regulated genes. The total number of DEGs is shown in circles and the numbers in between represent the overlapping set of genes (B-D). The significant *p-values* shown in red represent hypergeometric probability for the overlap. ns = not significant

**Fig. S15.**
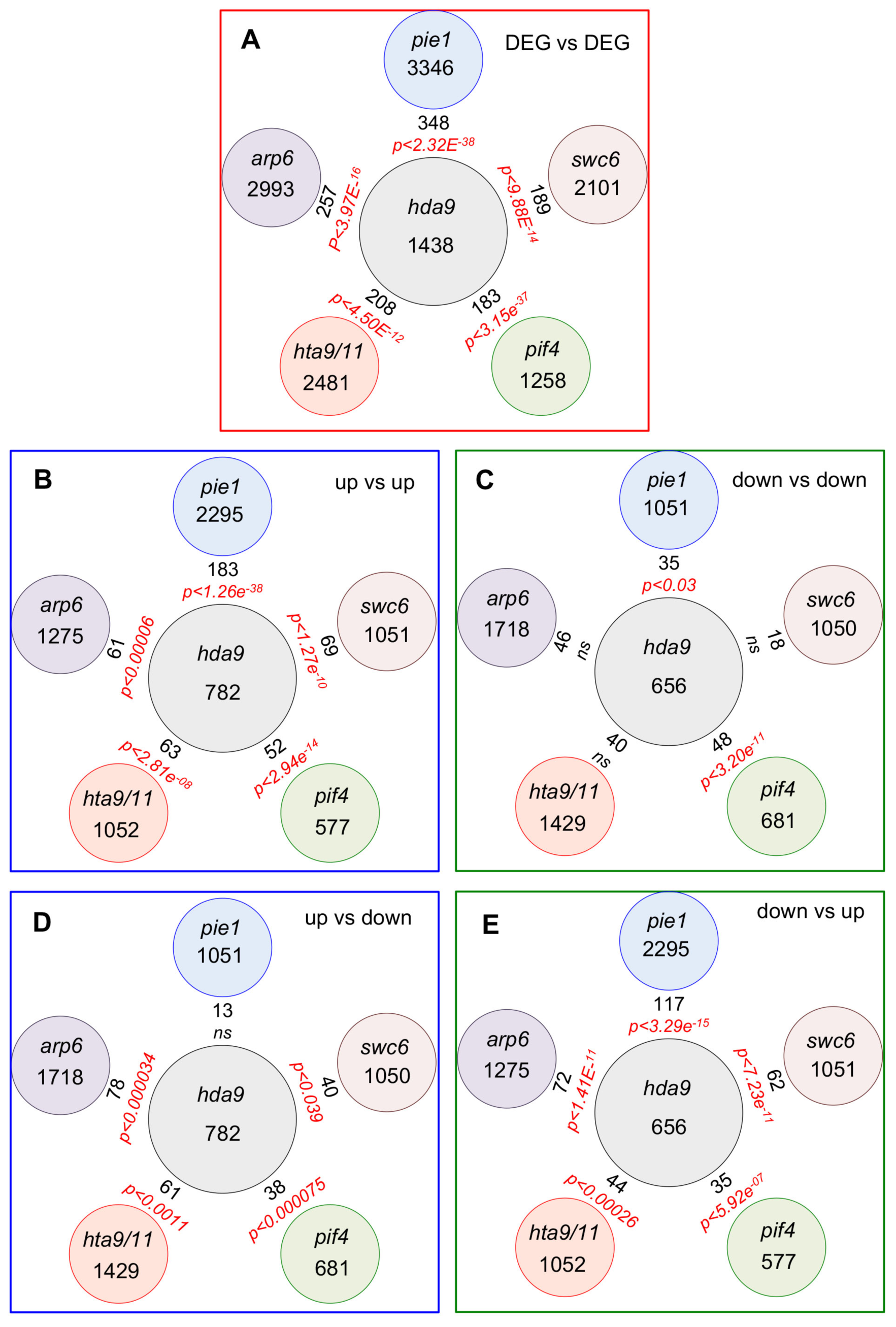
Overlap analysis of DEGs in *hda9* with DEGs in the transcriptomes of *pie1, swc6, arp6, pif4-2* and the *hta9/hta11* double mutants based on Chen et al data. A-E) Overlap of DEGs in *hda9* compared to Col-0 with DEGs in *pie1, swc6, pif4, hta9/hta11* and *arp6.* A) Overlap among DEGs. B) Overlap among genes that are up regulated in all genotypes. C) Overlap among genes that are down regulated in all genotypes. D) Overlap among genes that were up regulated in *hda9,* but down regulated in other genotypes. E) Overlap among genes that were down regulated in *hda9*, but up regulated in other genotypes. The total number of DEGs is shown in circles and the numbers in between represent the overlapping set of genes. The significant p-values shown in red represent hypergeometric probability for the overlap. ns = not significant The transcriptome data is from[18, 30, 33]

**Fig. S16.**
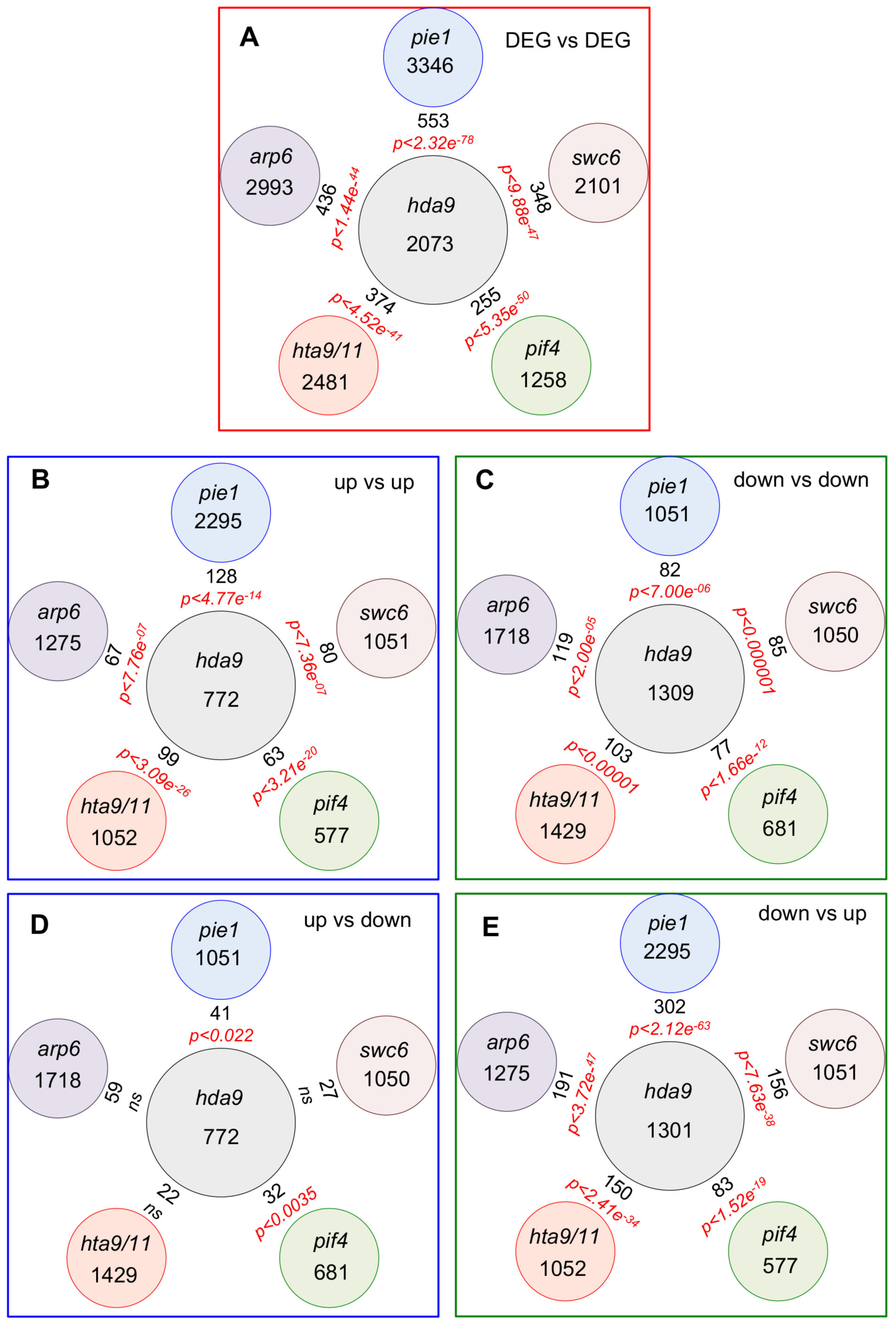
Overlap analysis of DEGs in *hda9* with DEGs in the transcriptomes of *pie1, swc6, arp6, pif4-2* and the *hta9/hta11* double mutants based on Kim et al data. A-E) Overlap of DEGs in *hda9* compared to Col-0 in the Kim et al data set with DEGs in *pie1, swc6, pif4, hta9/hta11* and *arp6.* A) Overlap among DEGs. B) Overlap among genes that are up regulated in all genotypes. C) Overlap among genes that are down regulated in all genotypes. D) Overlap among genes that were up regulated in *hda9,* but down regulated in other genotypes. E) Overlap among genes that were down regulated in *hda9*, but up regulated in other genotypes. The total number of DEGs is shown in circles and the numbers in between represent the overlapping set of genes. The significant p-values shown in red represent hypergeometric probability for the overlap. ns = not significant The transcriptome data is from[18, 20, 21, 30, 33]

**Fig. S17.**
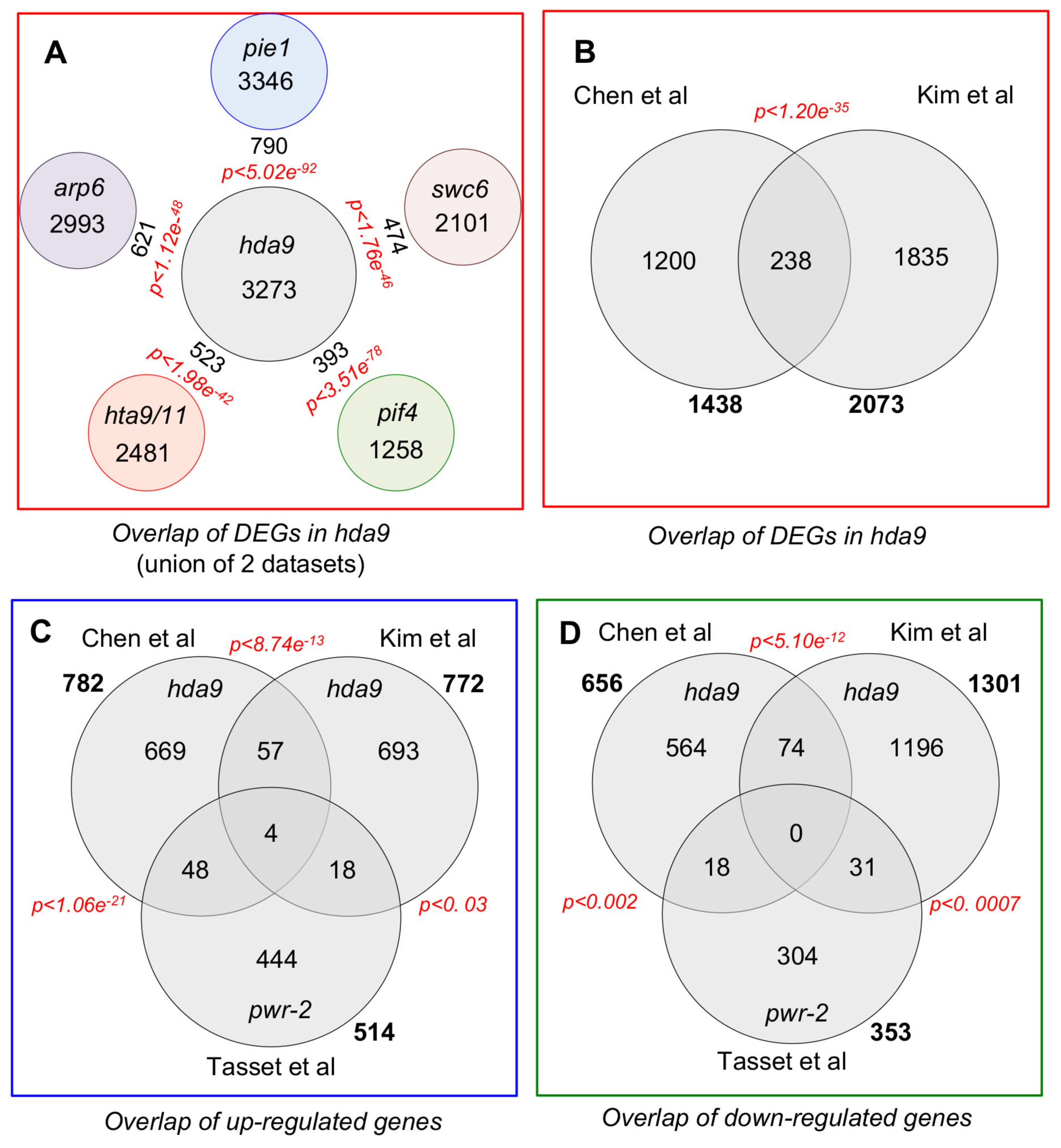
Comparison of the overlaps between the transcriptional response observed in *hda9* two different datasets and their overlap with the DEGs in *arp6, pie1, swc6, hta9/hta11* and *pif4.* A) Overlap of DEGs in *hda9* compared to Col-0 compiled from two datasets (excluding genes that did not change in the same direction in the datasets) with DEGs in *pie1, swc6, pif4, hta9/hta11* and *arp6.* The total number of DEGs is shown in circles and the numbers in between represent the overlapping set of genes. The transcriptome data is from [18, 20, 21, 30, 33]. B) Overlap of the DEGs in *pwr* in the three different datasets. The p-values are shown next to each of the overlaps. C) Overlap among up-regulated genes. D) Overlap among down regulated genes. The significant p-values shown in red represent hypergeometric probability for the overlap. ns = not significant.

**Fig. S18.**
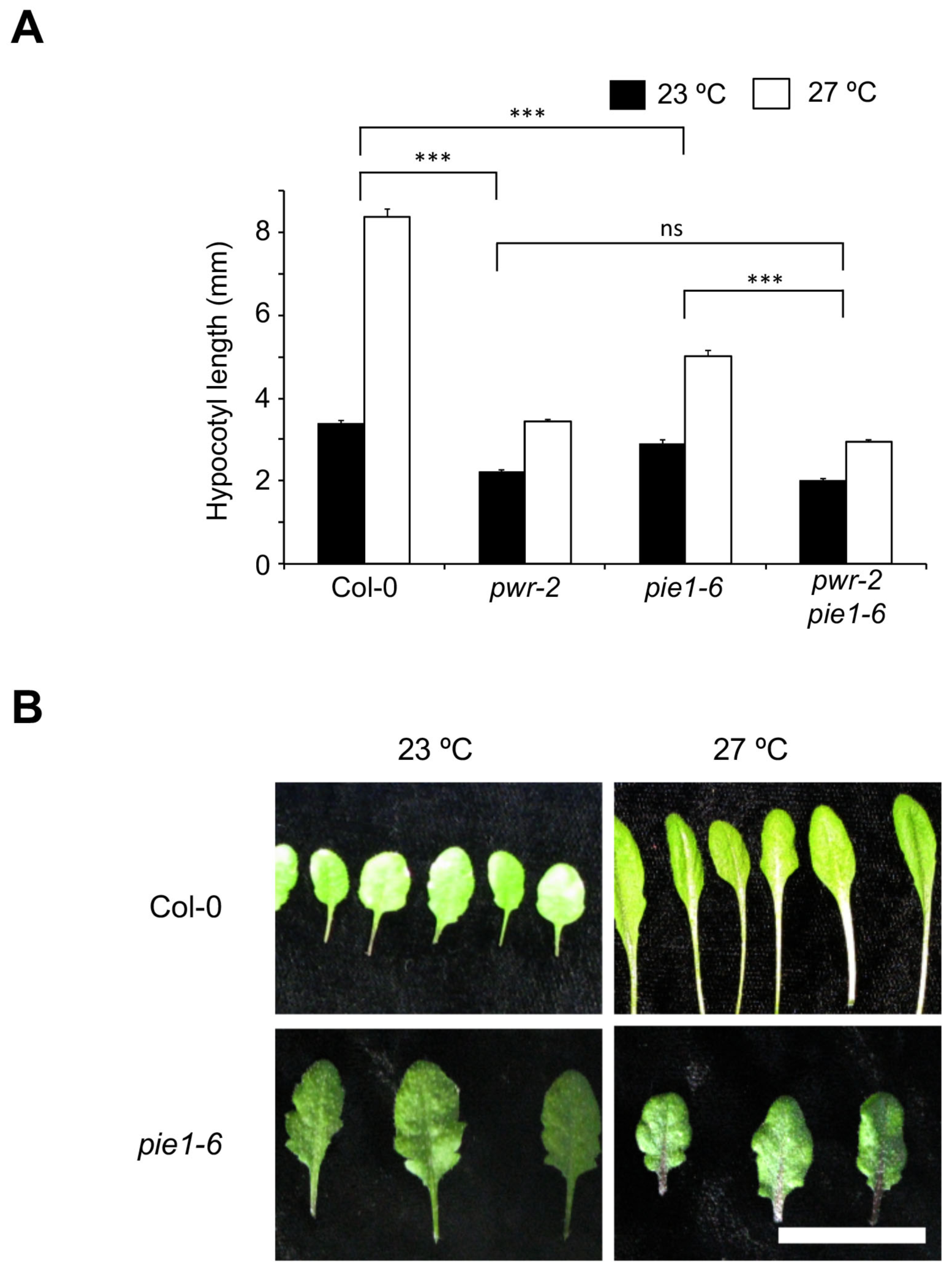
Mutations in *PIE1* compromise temperature-induced hypocotyl and petiole elongation. A) A) Hypocotyl lengths of various genotypes at 23 °C and 27 °C*. P-values* for the corresponding GxE interactions determined through ANOVA are shown. The Col-0 and *pwr-2* data is the same as shown in Fig 2A. B) Comparison of the Col-0 and *pie1* mutant leaves grown at 23 °C and 27 °C. Scale bar: 1cm. Error bars indicate standard error. *p‐ values*: ***<0.0001, **<0.001, *<0.05

## Supplementary Tables

**Table S1. List of DEGs between Col-0 and *pwr-2* at 23 °C**

**Table S2. List of DEGs between Col-0 and *pwr-2* at 27 °C.**

**Table S3. List of DEGs between 23 °C and 27 °C in Col-0**

**Table S4. GO-terms that were significantly enriched in DEGs in *pwr-2* mutants.** The GO terms with “response” are given in bold.

**Table S5. GO-terms that were significantly enriched in genes that were up-regulated in *pwr-2* mutants.** The GO terms with “response” are given in bold. The GO terms associated with “defense” are highlighted.

**Table S6. List of primers in used in this study.**

